# Bayesian efficient coding

**DOI:** 10.1101/178418

**Authors:** Il Memming Park, Jonathan W. Pillow

## Abstract

The efficient coding hypothesis, which proposes that neurons are optimized to maximize information about the environment, has provided a guiding theoretical framework for sensory and systems neuroscience. More recently, a theory known as the Bayesian Brain hypothesis has focused on the brain’s ability to integrate sensory and prior sources of information in order to perform Bayesian inference. Although pieces of a connection between these two hypotheses have appeared in prior work, a general formulation that treats the optimality criterion as an arbitrary functional of the posterior distribution – and thereby admits both information-based and other objectives within a single framework – has remained largely implicit. Here we make this formulation explicit, developing a Bayesian theory of efficient coding that defines Bayesian efficient codes in terms of four basic ingredients: (1) a stimulus prior distribution; (2) an encoding model; (3) a capacity constraint, specifying a neural resource limit; and (4) a loss functional, quantifying the desirability or undesirability of various posterior distributions. Classic efficient codes arise as the special case in which the loss functional is the posterior entropy, leading to a code that maximizes mutual information, but alternate loss functionals give solutions that differ dramatically from information-maximizing codes. Within this framework we introduce *covtropy*, a novel family of losses defined on posterior distributions and parameterized by a single exponent, and use it to show that decorrelation of sensory inputs – optimal under classic efficient codes in low-noise settings – can be disadvantageous for objectives that penalize large errors. We then reanalyze Laughlin’s seminal data on contrast coding in the blowfly large monopolar cell, and find that the measured response nonlinearity is better explained by minimizing *L*_*p*_ reconstruction error with *p* = 1*/*2 than by information maximization, overturning a forty-year-old interpretation. Bayesian efficient coding thus enlarges the family of codes that are optimal under different objectives and provides a more general framework for understanding the design principles of sensory systems.

**Author summary:** Sensory neurons work under tight budgets. They represent a rich, noisy world with limited spikes, limited dynamic range, and limited energy. Two long-standing ideas address this problem. One, called efficient coding, asks *why* neurons encode signals in the ways they do: which signals to amplify, which to filter out. A second, the Bayesian brain hypothesis, asks *how* the brain turns those signals into a percept: how to combine sensory evidence with prior knowledge to form a best guess about the world. We develop a framework that asks both questions at once. An optimal sensory code in our framework is specified by four interlocking ingredients: a prior description of the world, a model of how neurons respond to it, a limit on the resources they can spend, and a rule for what counts as a good internal representation. Classical information-maximizing codes emerge as one special case, one corner of a much larger family of Bayesian efficient codes. Applying this framework to Laughlin’s 1981 blowfly experiment, long held up as the textbook example of information maximization, we find that the fly’s neurons are better explained by minimizing decoding error than by maximizing information.

## Introduction

One of the primary goals of theoretical neuroscience is to understand the functional organization of neurons in the early sensory pathways and the principles governing them. Why do sensory neurons amplify some signals and filter out others? What can explain the particular configurations and types of neurons found in early sensory system? What general principles can explain the solutions evolution has selected for extracting signals from the sensory environment?

Two of the most influential theories for addressing these questions are the “efficient coding” hypothesis and the “Bayesian brain” hypothesis. The efficient coding hypothesis, introduced by Attneave and Barlow more than seventy years ago, uses the ideas from Shannon’s information theory to formulate a theory of normatively optimal neural coding [1, 2]. The Bayesian brain hypothesis, on the other hand, focuses on the brain’s ability to perform Bayesian inference, and can be traced back to ideas from Helmholtz about optimal perceptual inference [3–7].

A substantial literature has sought to alter or expand the original efficient coding hypothesis [5, 8–20], and a large number of papers have considered optimal codes in the context of Bayesian inference [21–29]. Although this prior work has made important contributions, a general formulation that treats the loss as an arbitrary functional of the posterior distribution – and thereby admits both information-theoretic and non-information-theoretic optimality criteria within a single formalism – has remained largely implicit. Here we make this formulation explicit, developing a general Bayesian theory of efficient coding that connects the two hypotheses. We begin by reviewing the key elements of each theory and then describe a framework for unifying them. Our approach involves combining a prior and model-based likelihood function with a neural resource constraint and a loss functional that quantifies what makes for a “good” posterior distribution. We show that classic efficient codes arise when we use information-theoretic quantities for these ingredients, but that a much larger family of Bayesian efficient codes can be constructed by allowing these ingredients to vary. We explore Bayesian efficient codes for several important cases of interest, namely linear receptive fields and nonlinear response functions. The latter case was examined in an influential paper by Laughlin that examined contrast coding in the blowfly large monopolar cells (LMCs) [30]; we reanalyze data from this paper and argue that LMC responses are better explained by minimizing *L*_*p*_ reconstruction error with *p* = 1*/*2 than by infomax.

## Theoretical background

### Efficient coding hypothesis

The Efficient Coding hypothesis, set forth by Attneave [1] and formalized by Barlow [2], proposes that sensory neurons are optimized to maximize the information they transmit about sensory inputs. This hypothesis represents one of the most influential theories in systems neuroscience, and was the first to apply Shannon’s information theory to the problem of neural coding.

Mutual information, as defined by Shannon [31], quantifies (in units of bits) the information between an external stimulus **x** and a neural response **y**, defined as *I*(**x, y**) = *H*(**y**) − *H*(**y**|**x**), where *H*(**y**) = − *∫ P* (**y**) log *P* (**y**) d**y** is the marginal entropy and *H*(**y**|**x**) = − ∫ *P* (**x**) ∫ *P* (**y**|**x**) log *P* (**y**|**x**) d**y** d**x** is the conditional (or “noise”) entropy of the response. The marginal entropy quantifies the uncertainty of the marginal response distribution *P* (**y**), while conditional entropy quantifies the uncertainty of the conditional response distribution *P* (**y**|**x**), averaged over the stimulus distribution *P* (**x**). The mutual information tells us how much uncertainty about **y** is reduced by knowing the stimulus **x**, on average. Remarkably, the mutual information is symmetric, so it can equally be written as *H*(**x**) − *H*(**x**|**y**), the difference between stimulus entropy *H*(**x**) and conditional entropy *H*(**x**|**y**), corresponding to the average reduction in uncertainty about the stimulus due to an observed neural response [32].

Barlow’s proposal, which he termed the *redundancy-reduction hypothesis*, was that neurons maximize *I*(**x, y**)*/C*, the ratio between the mutual information between stimulus **x** and neural response **y**, and the channel capacity *C*, which is an upper bound (considered over all stimulus distributions) on the mutual information. A perfectly efficient code in Barlow’s sense is one for which *I*(**x, y**) = *C*, where mutual information equals the channel capacity.

Barlow’s original work focused on the deterministic or ‘noiseless’ case where noise entropy is zero and the channel capacity is fixed. In this setting, efficiency is achieved by maximizing response entropy *H*(**y**). This specific setting yields two predictions most commonly associated with the efficient coding hypothesis: (1) single neurons should nonlinearly transform the stimulus to achieve optimal use of their full dynamic range [30, 33, 34]; and (2) neural populations should decorrelate their inputs, so the population responses are marginally independent [9, 35–38]. Subsequent work by Atick & Redlich [10, 39, 40] extended this framework to noisy responses, showing that information maximization predicts linear receptive fields that are low-pass at low light levels, where signal-to-noise is low, and band-pass or “whitening” at high light levels, where signal-to-noise is high, consistent with receptive field measurements in retina.

### Bayesian brain hypothesis

The Bayesian Brain hypothesis provides a second theoretical perspective on neural coding [4, 6, 7]. The core idea can be traced back to Helmholtz’s 19th century work on perception: it proposes that the brain seeks to combine noisy sensory information about the current stimulus with prior information about the environment. The product of these two terms, known as likelihood and prior, can be normalized to obtain the posterior distribution, which captures all information about the state of the environment. In this view, sensory perception is a form of Bayesian inference, and the brain’s goal is to compute the posterior distribution over environmental variables given sensory inputs.

The two key ingredients for the Bayesian Brain hypothesis are a prior distribution *P* (**x**) over the stimulus and an encoding distribution *P* (**y**|**x**) that describes the mapping from stimuli **x** to neural responses **y**. These ingredients combine according to Bayes’ rule to form the posterior distribution: *P* (**x**|**y**) ∝ *P* (**x**)*P* (**y**|**x**), which captures the observer’s beliefs about the stimulus **x** given the noisy sensory information contained in **y**. This theory has had major influences on the study of sensory and motor behavior [6, 7, 28, 41–47], as well as on theories of neural population codes that support Bayesian inference [21–23, 48–50]. Despite its emphasis on normatively optimal perception and behavior, the Bayesian brain literature and the efficient coding hypothesis have not yet been connected in full generality.

## Bayesian efficient coding

Here we make the connection between the Bayesian brain hypothesis and efficient coding explicit by formulating a more general theory of Bayesian efficient coding, which includes classic efficient coding as a special case. We will define a Bayesian efficient code (BEC) with four basic ingredients:

1. *P* (**x**) - A stimulus distribution, or prior.
2. *P* (**y**|**x**, *θ*) - An encoding model, parametrized by *θ*, describing how stimuli **x** are mapped to responses **y**.
3. *C*(*θ*) - A capacity constraint on the parameters, specifying a neural resource limit.
4. *L*(·) - A loss functional, quantifying the desirability or undesirability of various posterior distributions.

Given these ingredients, a BEC corresponds to the model parameters *θ* that achieve a minimum of the expected loss,

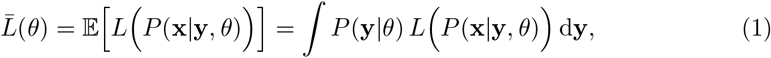

subject to the capacity constraint *C*(*θ*) ≤ **c**, where *P* (**x**|**y**, *θ*) ∝ *P* (**y**|**x**, *θ*)*P* (**x**) denotes the posterior over **x** given **y** and *θ*, and expectation is taken with respect to the marginal response distribution *P* (**y**|*θ*) = ∫ *P* (**y**|**x**, *θ*)*P* (**x**)d**x**.

Each of the four ingredients plays a distinct role in determining a Bayesian efficient code (see Fig. 1). The prior is determined by the statistics of the environment, and provides a complete description of the stimuli to be encoded. The model, in turn, determines the form of the probabilistic encoding function to be optimized; it determines the posterior distributions that the organism will utilize when the prior and likelihood are combined after each sensory response **y**^∗^, where the asterisk denotes a particular observed response.

**Figure 1.**
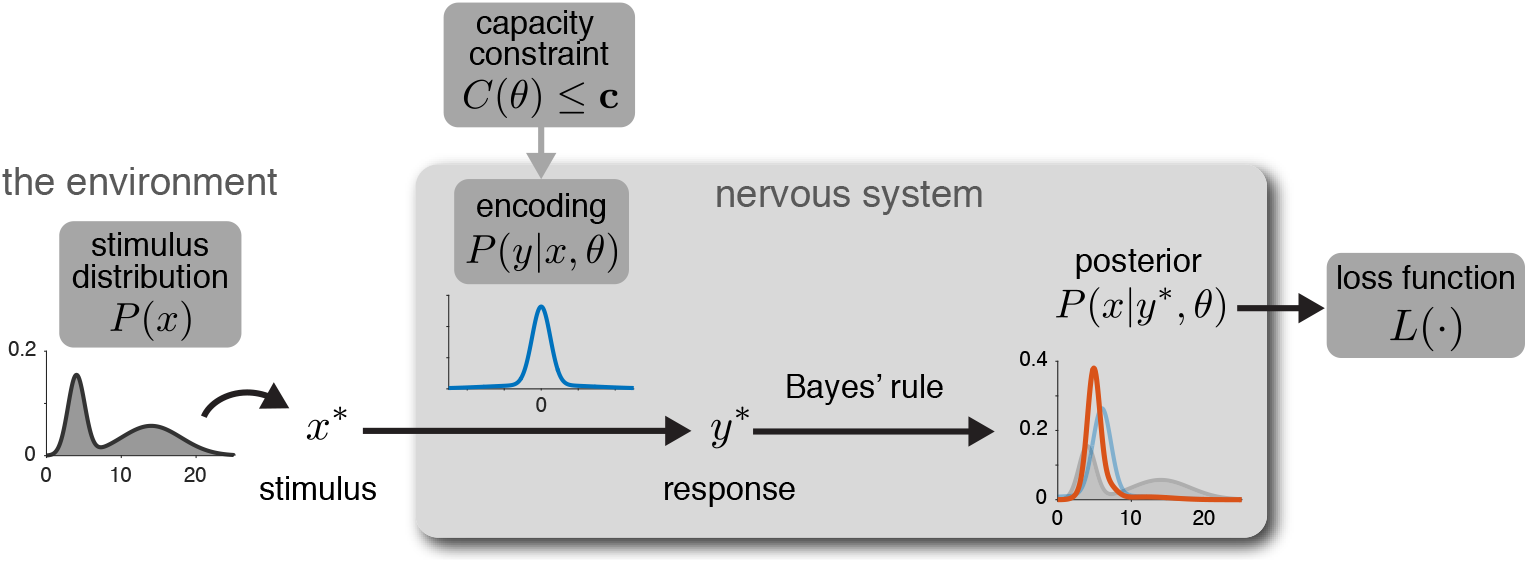
Bayesian efficient coding schematic. The theory is governed by four basic ingredients, highlighted in dark gray boxes. During any perceptual interval, a stimulus **x**^∗^ is drawn from the prior *P* (**x**) and presented to the organism. The nervous system encodes this stimulus with a sample **y**^∗^ from the encoding distribution *P* (**y**|**x**^∗^, *θ*), which is governed by parameters *θ* (e.g., defining a neuron’s receptive field, nonlinearity, noise, etc). An application of Bayes’ rule leads to the posterior distribution *P* (**x**|**y**^∗^, *θ*), which captures all information available to downstream brain areas about the stimulus given the sensory response **y**^∗^. The loss function *L*(·) characterizes the desirability of this posterior. A Bayesian efficient code is one for which parameters *θ* are set to minimize the average loss over stimuli and responses drawn from the prior and encoding model, subject to the resource constraint on the encoder, *C*(*θ*) *<* **c**.

The capacity constraint *C* defines a neural resource limitation, such as an energetic constraint on the average spike count or a physiological limit on the maximal firing rate. This makes the problem of determining an optimal code well posed, since without a constraint it is often possible to achieve arbitrarily good codes, e.g., by using arbitrarily large spike counts to encode stimuli with arbitrarily high fidelity.

Finally, the loss functional *L* is the key component that sets Bayesian efficient codes apart from classic efficient codes. It quantifies how much to penalize different posterior distributions that may arise from different settings of the encoder. In essence, the loss functional determines what counts as a “good” posterior distribution. For example: is it preferable for an organism to have a posterior distribution with low entropy, low variance, or low standard deviation? These desiderata are not the same, as we shall see shortly. We will discuss motivations for different choices of loss in the following sections.

### Bayesian vs. classical efficient codes

Given the above definition, it is natural to ask: when does a Bayesian efficient code correspond to a traditional efficient code as defined by Barlow? The answer is that classical Barlow efficient codes are a special case of Bayesian efficient codes with loss function set to the posterior entropy:

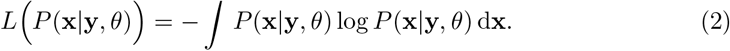

This results in a code that maximizes mutual information between stimulus and response because the expected loss 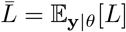 is the conditional entropy of the stimulus given the response *H*(**x**|**y**; *θ*); subtracting this from the prior entropy *H*(**x**), which is independent of *θ*, gives the mutual information *I*(**x, y**). Thus, minimizing average posterior entropy is the same as maximizing mutual information.

However, there is nothing privileged about minimizing entropy or maximizing information from a general coding perspective. In the following sections, we will show that it is often natural to consider other loss functions, and that the Bayesian efficient codes that result from doing so can differ strongly from classical, information-theoretically optimal codes.

## Simple examples

We will motivate a Bayesian theory of efficient coding by showing two simple examples that illustrate the appeal of using loss functions other than posterior entropy, one with continuous and one with discrete encoding.

### Continuous example: 2D Gaussian

To illustrate the role played by the loss function in Bayesian efficient codes, we first consider a toy example with two neurons encoding a bivariate Gaussian stimulus (Fig. 2A). Suppose that a 2D stimulus **x** = (*x*_1_, *x*_2_) has an independent Gaussian prior distribution with standard deviation *σ*_prior_ = 10 in both directions. We then consider three arbitrarily selected noisy encoders, each defined by a different level of independent additive Gaussian noise along each dimension:

**Figure 2.**
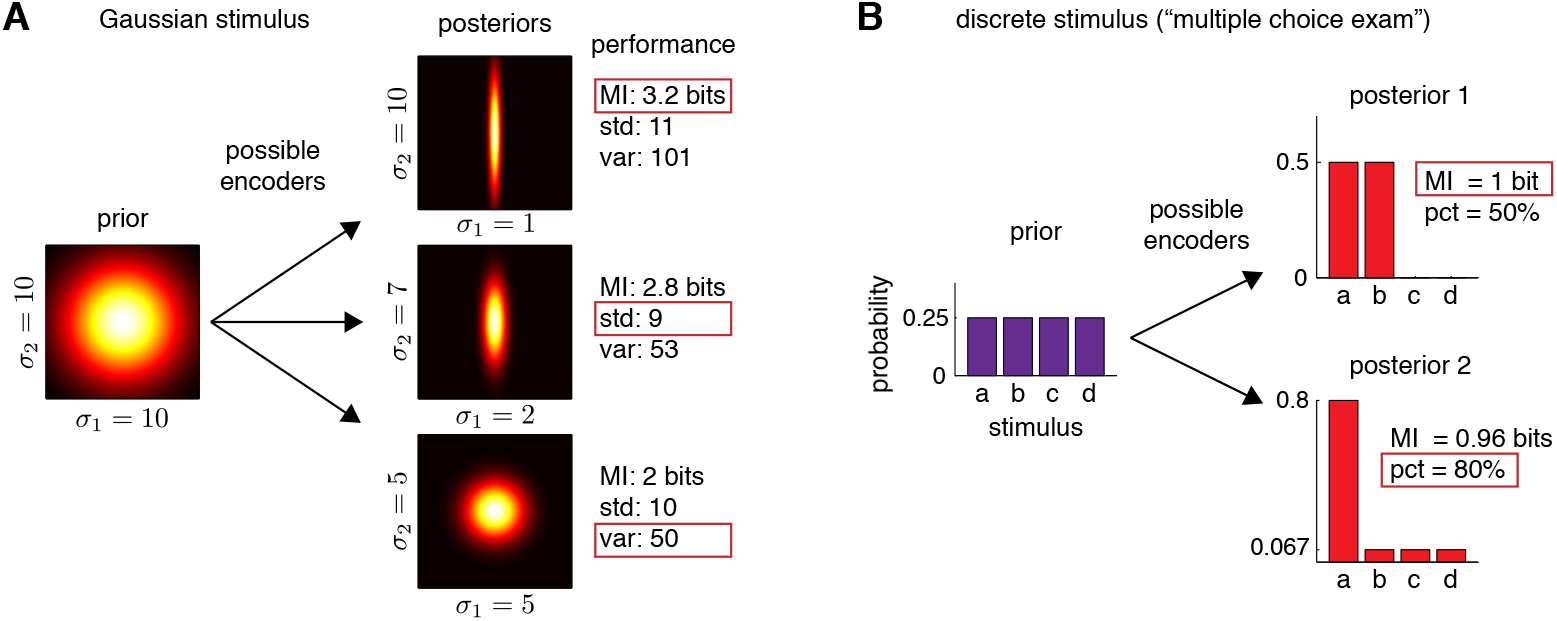
Optimal encoding depends critically on choice of loss function. **(A)** Illustration showing three possible encoders of a stimulus with an independent bivariate Gaussian prior distribution. The encoders produce three different Gaussian posteriors, shown at right. For axis-aligned Gaussian distributions, the entropy depends on the product of standard deviations *σ*_1_*σ*_2_, whereas the “total deviation” depends on the sum of standard deviations *σ*_1_ + *σ*_2_, and total variance depends on the sum of variances 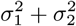. According to entropy loss, the top encoder is best (achieving a mutual information of log_2_(10 · 10) − log_2_(10 · 1) = 3.2 bits), but for total-deviation loss, the middle encoder is best (achieving *σ*_1_ + *σ*_2_ = 9), and for total-variance loss, the bottom encoder is best (achieving 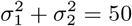). **(B)** Discrete encoding example, showing two possible encoders for a multiple-choice exam. The prior is an equal *p* = 1*/*4 probability for each of the four possible stimuli *{a, b, c, d}*. The top encoder eliminates two possibilities so that the posterior assigns probability *p*_*i*_ = 0.5 to two stimuli and 0 to the other two. The bottom encoder gives a posterior distribution with probability *p*_*i*_ = 0.8 for one stimulus and the remaining 0.2 probability spread evenly among the other three. The top encoder is optimal for information theoretic loss, since it achieves a mutual information of 1 bit, vs. only 0.96 bits for the bottom encoder. However, for “percent correct” loss, which is sensitive only to max(*{p*_*i*_*}*), the bottom encoder is clearly better. It identifies the correct stimulus 80% of the time (good for a grade of B-), whereas the top encoder achieves only 50% (an F).

- **encoder 1**: High-fidelity (i.e., low-noise) encoding of dimension *x*_1_ and zero-fidelity (i.e., infinite-noise) encoding of *x*_2_, resulting in a posterior with standard deviations (*σ*_1_, *σ*_2_) = (1, 10).
- **encoder 2**: Medium-fidelity encoding of dimension *x*_1_ and low-fidelity encoding of *x*_2_, resulting in posterior standard deviations (*σ*_1_, *σ*_2_) = (2, 7).
- **encoder 3**: Equal-fidelity encoding of both dimensions, resulting in posterior standard deviations (*σ*_1_, *σ*_2_) = (5, 5).

See Fig. 2A for an illustration. (See Methods for details).

It should be obvious that there is no loss-independent sense in which one of these three posteriors is preferable to the others. The three posteriors differ in how uncertainty is distributed across the two stimulus dimensions; which is better depends entirely on what the organism cares about. To make this concrete, we consider three loss functions: the posterior entropy, *L*_1_ = log(2*πe σ*_1_*σ*_2_), (equivalent to maximizing mutual information), the total standard deviation, *L*_2_ = *σ*_1_ + *σ*_2_, and the total variance, 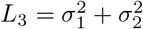. Each loss function gives a different best encoder. The first encoder achieves the smallest entropy (i.e., smallest product of standard deviations, 1 · 10 = 10, compared to 2 · 7 = 14 for encoder 2, or 5 · 5 = 25 for encoder 3); it thus achieves the highest mutual information, even though it throws away all information about *x*_2_. The second encoder achieves minimal total-deviation loss, because 2 + 7 = 9 (encoder 2) is less than 1 + 10 = 11 (encoder 1) or 5 + 5 = 10 (encoder 3). The third encoder minimizes total-variance loss *L*_3_, because 25 + 25 = 50 is smaller than either 1 + 100 = 101 (encoder 1) or 4 + 49 = 53 (encoder 2).

This simple example illustrates the manner in which different loss functions give rise to different notions of optimal coding. The loss functions differ in how they penalize the allocation of uncertainty across stimulus dimensions. Entropy is only sensitive to the product *σ*_1_*σ*_2_, which corresponds to the volume of the posterior. We could stretch the posterior by an arbitrary constant *a* in one dimension and by 1*/a* in the other, and we would not affect entropy. The other two loss functions, on the other hand, seek to minimize the summed uncertainty (standard deviation or variance) along the two axes. Compared to the entropy, they disfavor posteriors with large uncertainty in one direction, an effect that is larger for variance than for standard deviation. For Gaussian posteriors, they can also be interpreted as minimizing error in an optimal decoder.

Minimizing total standard deviation is equivalent to finding an encoder that minimizes the summed absolute error of a decoded estimate: 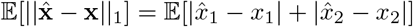, where 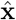 is an estimate that minimizes the *ℓ*_1_ error 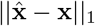. Similarly, minimizing total variance is equivalent to minimizing the mean squared error of an optimal decoder: 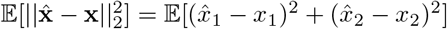, where 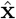 is the posterior mean (note that this connection between sum of moments and minimization of error does not hold in general; see S1 Appendix, Loss functionals), also known as the Bayes’ least squares (BLS) or minimum mean squared error estimator.

We could of course have considered other loss functions that would have given us different reasons for preferring any of these three encoders. For example, a “minimax” loss function *L* = max(*σ*_1_, *σ*_2_), which cares only about minimizing the worst possible performance in any dimension, would also favor the third encoder. Thus, optimality is in the eye of the loss function. For any set of encoders there may be multiple ways to regard them as Bayes optimal.

### Discrete example: multiple choice exam

As second example, consider the case of a discrete stimulus that takes on one of four possible values {*a, b, c, d}*, each with prior probability 0.25, and a noisy neuron that can respond with 1, 2, 3, or 4 spikes. We will consider the following two possible encoding rules (see Table 1 and Fig. 2B).

**Table 1.** Two encoding distributions *P* (*y*|*x*) for the multiple-choice exam example.

| $y \backslash x$ | $a$ | $b$ | $c$ | $d$ | $y \backslash x$ | $a$ | $b$ | $c$ | $d$ |
| --- | --- | --- | --- | --- | --- | --- | --- | --- | --- |
| 1 | 0.5 | 0.5 | 0 | 0 | 1 | 0.8 | 0.2/3 | 0.2/3 | 0.2/3 |
| 2 | 0.5 | 0.5 | 0 | 0 | 2 | 0.2/3 | 0.8 | 0.2/3 | 0.2/3 |
| 3 | 0 | 0 | 0.5 | 0.5 | 3 | 0.2/3 | 0.2/3 | 0.8 | 0.2/3 |
| 4 | 0 | 0 | 0.5 | 0.5 | 4 | 0.2/3 | 0.2/3 | 0.2/3 | 0.8 |

(a) encoder 1
(b) encoder 2

The first encoder maps stimuli *a* and *b* to responses 1 and 2 randomly with equal probability, and similarly maps stimuli *c* and *d* to responses 3 and 4. The second encoder, on the other hand, maps *a* → 1, *b* → 2, *c* → 3, *d* → 4 with probability 0.8, and with probability 0.2 maps to one of the other three responses selected at random. Fig. 2B shows the kinds of posteriors that arise under these two encoders—these are in fact equal to the rows of the encoding tables given above, since the prior is uniform and each row already sums to 1. For the first encoder, the posterior assigns probability 0.5 to two stimuli and rules out the other two. For the second encoder, the posterior concentrates on one stimulus with probability 0.8, and spreads the remaining 0.2 evenly across the other three stimuli.

It is now interesting to consider which of these two encoders is best according to different loss functions. First, let us consider posterior entropy, which corresponds to maximizing information as in Barlow’s classic definition. The prior has an entropy *H*(*x*) = −4 *×* 0.25 log_2_ 0.25 = 2 bits. The mean posterior entropy of the first encoder *H*(*x*|*y, θ*_1_) = −2 *×* 0.5 log_2_ 0.5 = 1 bit, so the mutual information between stimulus and response is 1 bit. By contrast, the posterior entropy of the second encoder is *H*(*x*|*y, θ*_2_) = − 0.8 log_2_ 0.8 − 0.2 log_2_(0.2*/*3) = 1.04 bits, which gives mutual information of only 2 − 1.04 = 0.96 bits. This means that the second encoder preserves strictly less Shannon information about the stimulus than the first, so the first encoder is more efficient (strictly speaking, the two channels have different capacity, so the definition of redundancy does not apply directly). A second natural choice of loss function, however, is the “percent correct”, defined as the percent of the time a *maximum a posteriori* decoder chooses the correct stimulus. According to this loss function, the second encoder is clearly superior, since 80% of the time it concentrates on the correct stimulus, whereas the best possible decoding of responses from the first encoder can only answer correctly 50% of the time.

This example has perhaps greater cultural and psychological salience if we reframe it terms of students studying for a multiple choice exam. Each question on the exam will have four possible choices (*a, b, c*, and *d*), which occur equally often. Student 1 adopts a study strategy that allows her to rule out two of the four choices with absolute certainty, but to have total uncertainty about which of the two remaining options is correct for each exam question. Student 2, on the other hand, adopts a study strategy that allows her to know the correct answer 80% of the time, but has uniform uncertainty about the remaining three options, which are correct the remaining 20% of the time. Which student’s strategy is better? If we judge them according to mutual information, the first student has clearly learned more; her brain has stored 0.04 more bits about the subject matter than the second student. However, if we judge them according to the number of questions they can answer correctly on the exam, the second student’s strategy is clearly better: her expected grade is a B-, with a score of 80%, whereas the first student is expected to fail with only half the questions answered correctly.

An interesting corollary of this example, therefore, is that although information-theoretic learning (cf. [12, 51]) is optimal for True/False exams, it can be substantially sub-optimal for multiple-choice exams.

## Linear receptive fields

Now we turn to an application motivated by biology. Neurons in the early visual system are often described as performing an approximately linear transformation of the light pattern falling on the retina. A large body of previous work has examined the optimality of these linear weighting functions under the efficient coding paradigm and its variants [10, 15, 16, 18, 36, 39, 40, 52–55].

Here we re-examine this problem through the lens of Bayesian efficient coding. We consider the following simplified model for linear encoding of sensory stimuli:

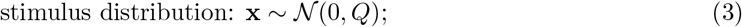

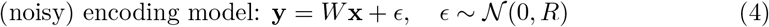

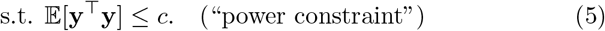

In this setup, we assume that the stimulus, an image consisting of *n* pixels, is encoded into a population of *n* neurons via a linear weight matrix *W*. Each row of *W* corresponds to the linear receptive field of a single neuron. For tractability, we take the population size equal to the stimulus dimensionality, which supplies the shared eigenbasis that makes the problem analytically solvable, and we assume that the stimulus has a Gaussian distribution (with covariance *Q* defining the correlations between pixels), and the neural population response is corrupted by additive Gaussian noise with covariance *R*. We impose a power constraint on the response, corresponding to a bound on the sum of squared responses **y**. This is a common choice of constraint in the efficient coding and engineering literature due to differentiability and analytic tractability [32, 39]. In practice, the power constraint restricts the model from achieving infinite signal-to-noise ratio by growing *W* without bound.

For this model, the marginal response distribution *P* (**y**) is Gaussian, **y** ~ *N* (0, *WQW*^⊤^ + *R*), so the power constraint can be written in terms of the trace of the marginal covariance:

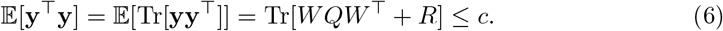

The posterior distribution *P* (**x**|**y**) is also Gaussian, *N* (*µ*, Σ), with mean and covariance

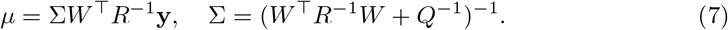

In this setup, the coding question of interest is: given stimulus covariance *Q*, noise covariance *R*, and power constraint *c*, what is the optimal linear encoding matrix *W* ? That is, what receptive fields are optimal for encoding stimuli from *P* (*x*) in the face of noise and a constraint on response variance? Here we consider two possible forms for the loss functional:

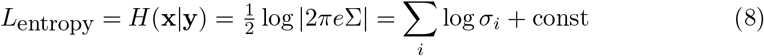

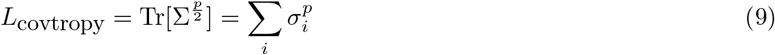

where 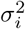 are the eigenvalues of posterior covariance Σ, or the variances of the posterior along its principal axes.

The first loss function, posterior entropy, corresponds to classic *infomax* efficient coding, since minimizing posterior entropy corresponds to maximizing mutual information between stimulus and response. The second loss function, which we introduce here and term *covtropy* due to its similarity to entropy, is the summed *p*’th powers of posterior standard deviation along each principal axis. Covtropy is, to our knowledge, a novel family of posterior-functional loss functions, parameterized by a single exponent *p*, that interpolates between infomax coding (*p* → 0), mean-squared-error coding (*p* = 2), and mini-max coding (*p* → ∞), and is rotation-invariant in stimulus space. Like entropy, the covtropy depends only on the eigenvalues of the posterior covariance matrix, and is thus invariant to rotations (see S1 Appendix, Loss functionals). For *p* = 2, covtropy corresponds to the sum of posterior variances along each axis; minimizing it is equivalent to minimizing the mean squared error [15–17] (see S1 Appendix, Equivalence of MSE and covtropy for *p* = 2 for proof).

For *p* = 1, covtropy corresponds to a sum of standard deviations; this penalizes a posterior with large variance in one direction less severely than covtropy with *p* = 2. In the limit *p* → 0, minimizing covtropy is identical to maximizing mutual information for Gaussian posteriors, since 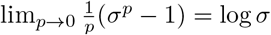. In this limit, the optimal code minimizes the sum of log standard deviations, ∑_*i*_ log *σ*_*i*_, which is equivalent to minimizing the product Π_*i*_ *σ*_*i*_. (Note that this equivalence relies on the posterior entropy being a function of the covariance alone, which holds for Gaussian but not general distributions; see S1 Appendix, Loss functionals.) The other interesting regime to consider is the limit *p* → ∞. In this regime, minimizing covtropy is equivalent to minimizing max_*i*_ *{σ*_*i*_*}*, the maximal posterior standard deviation, a form of “mini-max” encoding. The optimal code is therefore the one that achieves the smallest maximal posterior standard deviation in any direction.

Note that for the linear Gaussian model considered here, the posterior covariance Σ is independent of the response **y**. This makes the problem easier to analyze because there is no need to compute average loss over the response distribution *P* (**y**) (eq. 1). We show here that there is an analytic solution for the optimal linear encoding matrix *W* in this setting, for both infomax loss and covtropy loss (with any choice of *p >* 0), if we assume that *Q, R*, and *W* have a common diagonalization. This condition arises naturally for a convolutional code, that is, *W* contains a single receptive field shape that is circularly shifted by one pixel for each neuron in the population, and noise is spatially shift invariant (i.e., *R* is a circulant matrix). In the following, we examine the properties of Bayesian efficient linear receptive field codes for both information-theoretic and covtropy loss functions (derivation in the SI Appendix).

### Infomax encoding

The optimal weight matrix *W* for infomax coding is given by:

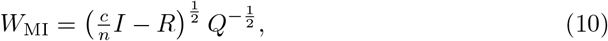

where *c* is the power constraint and *n* is the number of neurons. For this encoder, the responses **y** are perfectly whitened, meaning the response covariance is proportional to the identity, E[**yy**^⊤^] ∝ *I*, and the posterior covariance of **x**|**y** is proportional to the product of signal and noise covariances, Σ ∝ (*QR*).

### Minimum covtropy encoding

For covtropy loss with exponent *p*, the optimal encoding weights *W* are given by

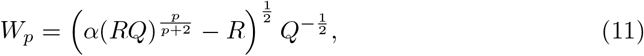

where constant 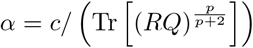 simply enforces the power constraint. The constrained solution used in the low-power regime is described in Methods. For the all-active weights, the marginal covariance is

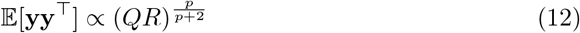

which implies that optimal responses can be more or less correlated than the stimulus. For example, when noise covariance *R* is proportional to stimulus covariance *Q*, then for *p* = 2 (minimum mean squared error encoding), the optimal receptive fields perfectly preserve the correlations in the stimulus, that is cov[*Y*] ∝ *Q*. For *p >* 2, the optimal receptive fields *increase* correlations so that responses are *more* correlated than the stimuli (See Fig. 3).

**Figure 3.**
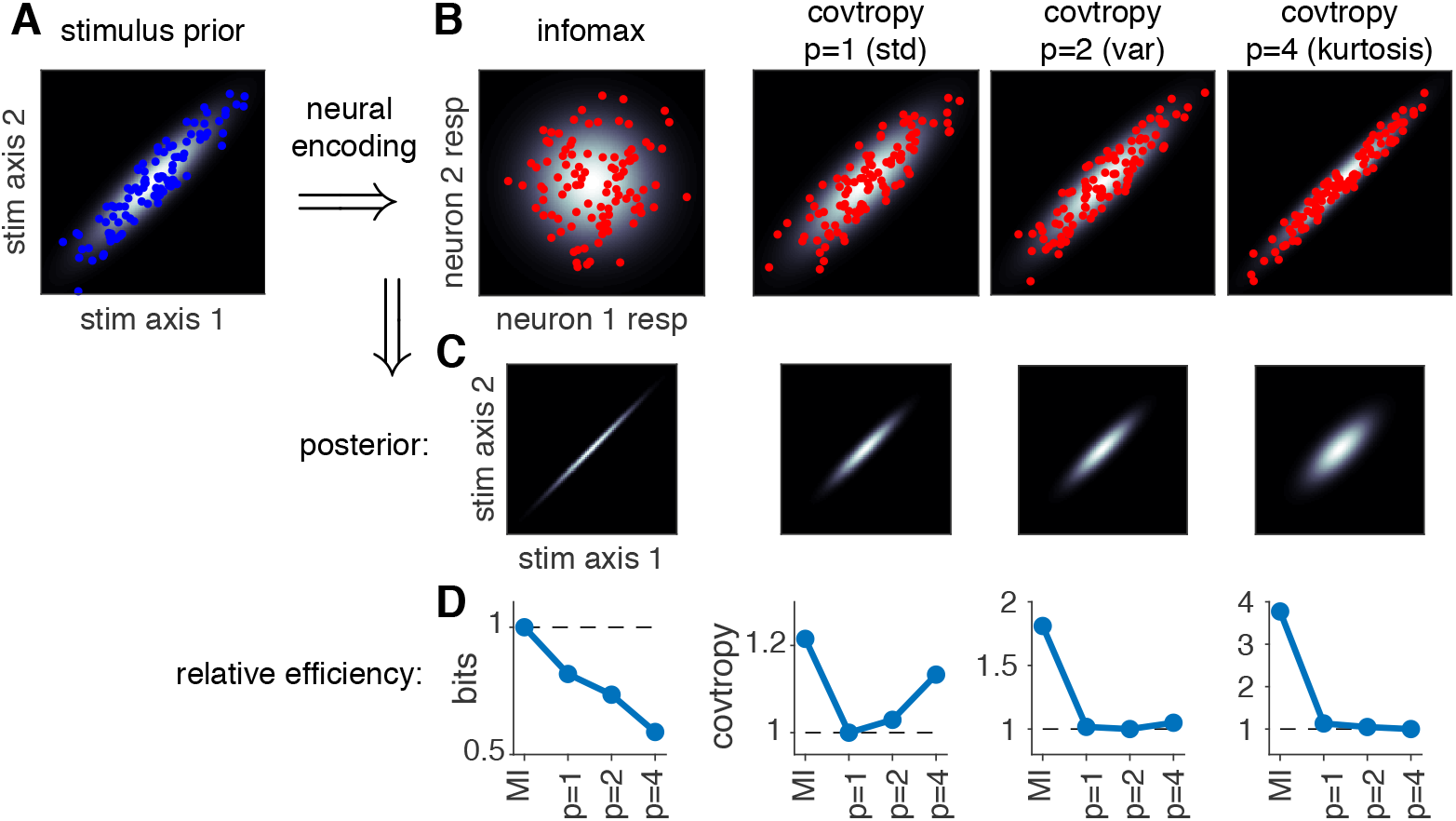
Bayesian efficient codes for a linear receptive field model with Gaussian noise. **(A)** Gaussian stimulus distribution with strong correlations (*ρ* = 0.9), with 50 samples shown (blue dots). **(B)** Neural responses (red dots) corresponding to optimal representations under infomax and covtropy loss functions, under infinitesimal Gaussian noise with covariance proportional to the stimulus distribution. Responses are uncorrelated (“white”) for the infomax encoder, but unchanged for the *p* = 2 (minimum variance) encoder and *stronger* for *p* = 4. **(C)** Posterior distribution shape for each optimal encoder, showing that the infomax encoder achieves small posterior entropy using a long and narrow posterior, whereas optimal covtropy posteriors grow more spherical with increasing *p*. **(D)** Relative performance of each encoder according to the loss function considered, normalized so the optimum is 1. Each encoder does best according to its own loss function, and the fall-off in efficiency between infomax and covtropy encoders is substantial; the infomax coder captures almost twice as much information as the *p* = 4 covtropy encoder in terms of bits (left plot), while the infomax coder exhibits nearly 4 times higher *p* = 4 covtropy than the optimal *p* = 4 encoder (right plot).

Although the responses become more correlated with increasing *p*, the posterior covariance, given by

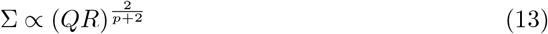

becomes increasingly white (i.e., uncorrelated) with increasing *p*, due to the fact that lim_*a*→0_ *M*^*a*^ = *I* for any nonsingular matrix *M*. This results in a posterior with the same uncertainty in all directions, regardless of the prior or noise covariance, in accordance with the “minimax” property for *p* = ∞ noted above. Thus, for example, if the additive noise is isotropic, the optimal filter bank in the *p*→ ∞ limit will be the identity matrix, regardless of the prior.

Fig. 4 shows the shape of the optimal high-dimensional linear RF for encoding stimuli with naturalistic (1*/f* ^2^) power spectrum in a population of neurons with i.i.d. Gaussian noise. Even in the high-SNR regime (when noise is infinitesimal), where perfect whitening maximizes mutual information, the optimal RFs under covtropy loss give less whitening as a function of increasing covtropy exponent *p*.

**Figure 4.**
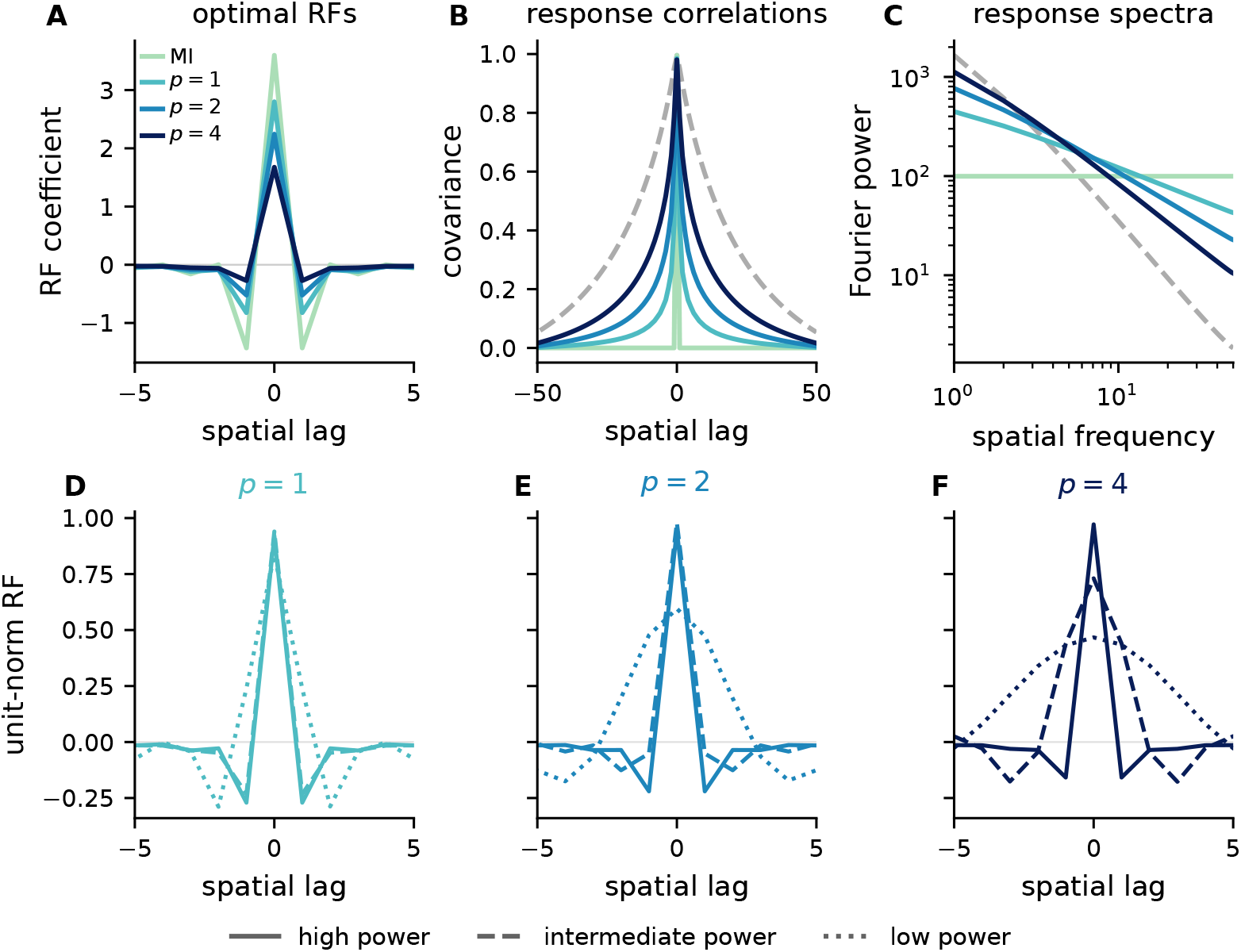
Optimal linear receptive fields for different loss functions and power constraints. **(A)** Optimal 1D receptive fields in raw coefficient units for a 100-neuron population encoding correlated Gaussian stimuli with a 1*/f* ^2^ power spectrum in the high-power regime. **(B-C)** Spatial covariance and Fourier power spectrum of the stimulus (dashed gray) and population responses. Infomax whitens the response, whereas covtropy encoders preserve more low-frequency information. **(D-F)** Unit-norm central RFs for *p* = 1, 2, 4 under high, intermediate, and low power constraints, with *Q* and *R* fixed. RF shape becomes more sensitive to the constraint as *p* increases. The infomax RF is omitted from D-F because its normalized shape is invariant to *c* for white output noise; exact parameter values are given in Linear receptive fields.

Holding *Q* and *R* fixed, we compared the high-power regime with progressively tighter response-power constraints (Fig. 4D-F). RF shapes are stable at high power and become increasingly sensitive to the constraint at low power, with larger changes for larger *p*. The qualitative ordering across loss functions nevertheless remains: larger *p* produces less whitening and greater preservation of low frequencies.

## Nonlinear response functions

So far we have focused on encoders that linearly transform the stimulus. However, many neurons exhibit rectifying or saturating nonlinearities that map the raw stimulus intensities so as to make optimal use of a neuron’s limited response range.

### Noiseless response

Barlow’s efficient coding theory states that the nonlinearity should map the stimulus distribution so as to maximize information between stimulus and response [2, 34], which naturally depends on the prior distribution over stimuli, the conditional response distribution, and the particular constraint on neural responses. For the case of noiseless responses and a simple constraint over the range of allowed responses, the shape of the information-maximizing nonlinearity is proportional to the cumulative distribution function (CDF) of the stimulus distribution, producing a uniform marginal distribution over responses [56].

In a groundbreaking paper that is widely considered the first experimental validation of Barlow’s theory, Simon Laughlin [30] measured graded responses from blowfly large monopolar cells (LMC) to contrast levels measured empirically in natural scenes. Laughlin found that the LMC response nonlinearity, measured with responses to sudden contrast increments and decrements, exhibited a striking resemblance to the shape of the empirically measured CDF of contrast levels in natural scenes. Fig. 5A shows a reproduction of the original figure, showing that the nonlinear response is closely matched with the stimulus distribution expected by the infomax solution.

**Figure 5.**
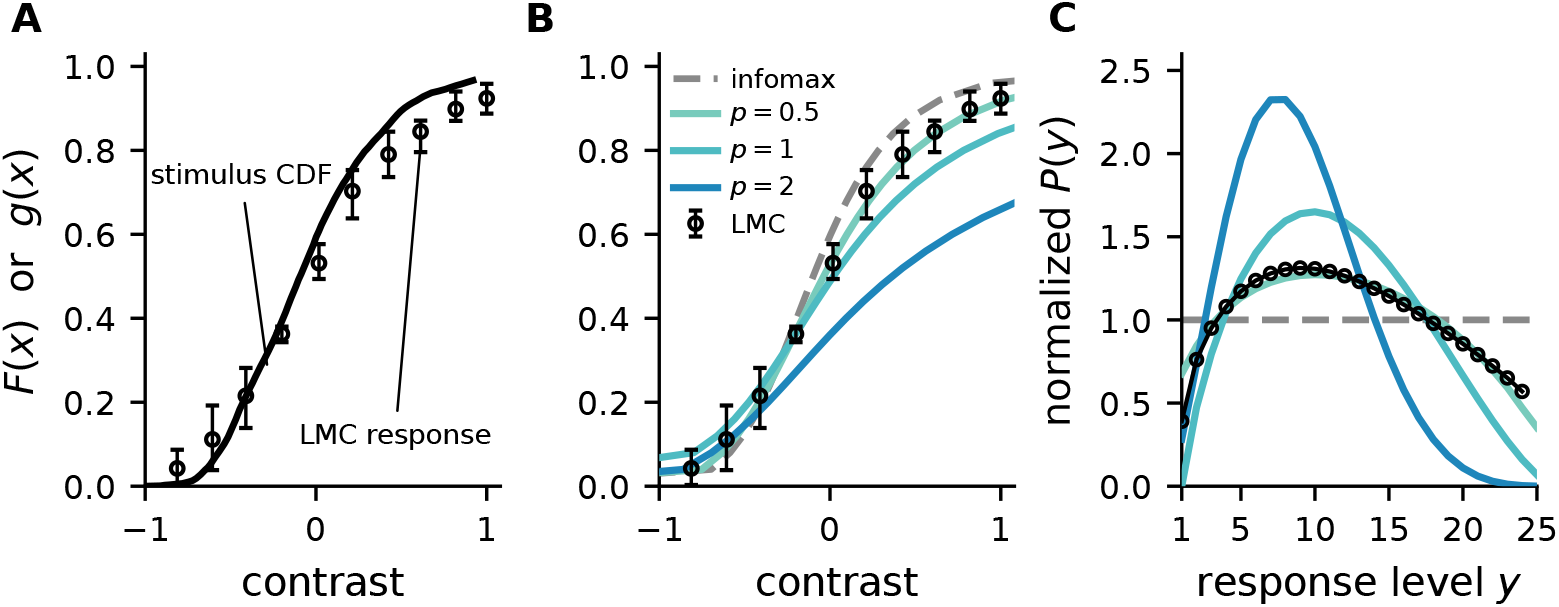
Bayesian efficient coding for noiseless nonlinear transformations applied to LMC response. **(A)**: Measured large monopolar cell (LMC) response from [30] (circles, mean *±* error bars) and the cumulative distribution function of the stimulus statistic (line). The two should be identical for optimal encoding under the infomax criterion. **(B)** Optimal nonlinearities predicted under *L*_*p*_ loss for 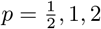, together with the infomax solution (*p* → 0, dashed), all computed from the Naka-Rushton fit of the stimulus distribution, against the same measured response. **(C)** Response marginals those codes induce, normalized so that infomax is flat at 1, and the marginal implied by the measured *g* (circles).

Here we re-examine Laughlin’s conclusions by analyzing the same dataset through the enlarged framework of Bayesian efficient coding. We can formalize the coding problem as:

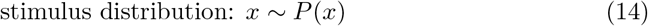

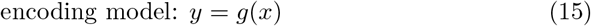

Here *x* is a scalar stimulus with prior distribution *P* (*x*) and the encoding model is described by *g*(*x*), a noiseless transformation from stimulus to discrete response levels. If we assume a finite *quantized* response *y* ∈ {0, …, *y*_*max*_*}*, some information about the stimulus is lost due to quantization error. For classical infomax encoding, we know that optimal *g* performs histogram equalization by taking on the quantiles of *P* (*x*), resulting in a uniform marginal response distribution *P* (*y*) [30, 56].

We consider several different choices of loss function, in particular, the *L*_*p*_-”norm” reconstruction error family of the form:

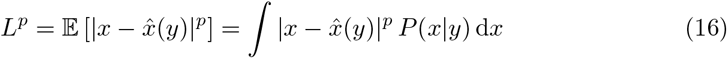

where *p* ≥ 0 and 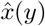 is the Bayesian optimal decoder for *x* from *y* [57]. For *p* = 2, this loss is equivalent to the mean squared error, and estimate 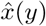 is equal to the mean of the posterior over *x* given *y*. Smaller values of *p* correspond to reducing the relative penalty on large errors and increasing the relative penalty on small errors; in the limit of *p* → 0, the loss converges to posterior entropy, making it equivalent to the infomax setting [26, 57].

We extracted the stimulus distribution from ([30], Fig. 2) to determine the optimal nonlinearity under Bayesian efficient coding paradigms. The cumulative stimulus distribution of the normalized contrast was fit with Naka-Rushton function which is a shifted log-logistic distribution (Fig. 5A). For this prior, the optimal solution can be derived in closed form (eq. 21, Closed-form optimal nonlinearity for a log-logistic prior) and it matches the numerically optimized quantization (Fig. S2).

The empirical nonlinearity (Fig. 5A) was also extracted from ([30], Fig. 2). We plotted the optimal nonlinearity *g*(*x*) family for *p* = {0.5, 1, 1.5, …, 4} (for details, see Fig. S3). We plotted these Bayes optimal nonlinearities against the infomax nonlinearity from Laughlin, as well as the empirically measured LMC response nonlinearity (Fig. 5B). We quantified agreement with the measured LMC response using the root-mean-square error (RMSE) across the ten digitized response points. Using the same fitted stimulus distribution, the RMSE decreases from 0.051 for infomax to 0.015 for BEC with 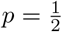, a reduction of approximately 70% (Fig. S3B). The 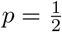 nonlinearity gives the lowest RMSE on the tested grid.

The difference between infomax and 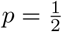 loss appears slight when viewed in terms of the optimal nonlinearity *g*(*x*), but is more dramatic when we consider the predicted marginal distribution over responses *P* (*y*) (Fig. 5C). We fit the neuron’s true response nonlinearity with a sigmoid function (Naka-Rushton CDF; See Methods), and computed the predicted marginal response distribution *P* (*y*) for each nonlinearity *g*(*x*). As expected, the infomax nonlinearity (dashed trace) produces a flat (uniform) distribution over response levels. However, both the 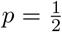 nonlinearity and the predicted LMC response distribution exhibit a noticeable peak around intermediate response levels, with intermediate response levels used more often, and responses at the extreme left and right tails used less often. We note that such peaking is expected under loss functions that penalize the magnitude of decoding errors. In fact, the optimal nonlinearities for *p* = 1 and *p* = 2 generate response distributions that are even more peaked than the real data, since they assign less probability mass to outermost response levels, where decoding errors are largest. The infomax loss function, by contrast, ignores the size of errors and simply seeks to match quantiles of the stimulus distribution.

Schaffner and colleagues [58] have since provided a task-level normative argument that supports the *p* = 1*/*2 result. They consider an organism performing repeated two-alternative forced choices over stimuli drawn from the natural distribution, with stimuli linearly mapped to reward value, and derive the utility function that maximizes expected reward without any assumption about maximizing information. The fitness-maximizing solution they obtain coincides with the *p* = 1*/*2 loss that we empirically identified here. Their derivation thus supplies an independent normative justification for the specific exponent that best fits the Laughlin 1981 LMC data, arrived at without invoking infomax as a goal in itself. For additional discussion of LMC response properties through a normative lens, see [59] (Chapter 8), which considers additional datasets from Laughlin [60] as well as the effects of temporal stimulus sequences [61].

### Linear-nonlinear-Poisson spiking

In the previous example, we assumed there was no variability in the neural response, but what if we extend the model to incorporate neural response noise? Here we examine optimal encoding in context of Poisson response noise to demonstrate the general principles of BEC. We consider the following linear-nonlinear-Poisson (LNP) model [62].

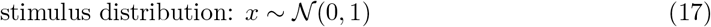

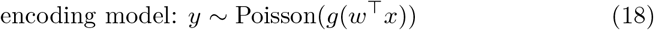

where *g*(·) ≥ 0 is a non-decreasing scalar function. We consider a constraint on the marginal firing rate, *C* = E[*y*] = *ĉ*, where *ĉ* is the empirical mean rate. Unlike the linear receptive field with Gaussian noise case discussed earlier, this form is no longer analytically tractable. Hence, we once again use numerical optimization to find the optimal nonlinearity *g* from neural spike counts.

We analyzed a blowfly H1 neuron’s responses to horizontal motion driven by white Gaussian noise [63]. We projected the stimulus onto its spike-triggered average (STA), then averaged the projections and counted spikes over 50 ms windows. Gaussian-process-Poisson regression provided the empirical nonlinearity and its uncertainty [64] (Fig. 6A). We compared this with non-decreasing encoders optimized under the *L*_*p*_-norm loss in eq. 16 and constrained to match the observed mean spike count (Fig. 6B).

**Figure 6.**
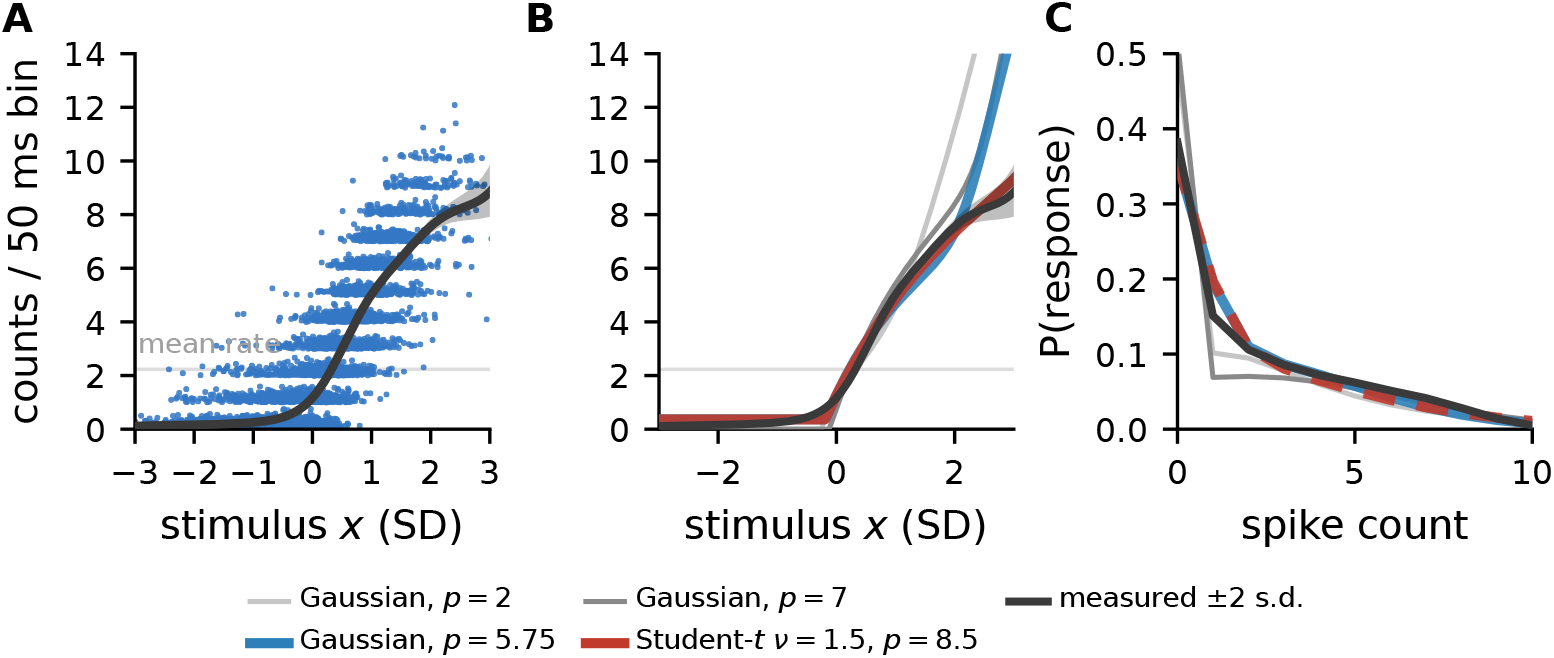
Bayesian efficient coding for linear-nonlinear-Poisson model applied to H1 response. **(A)** Blowfly H1 neuron response to white noise from [63]. Black curve shows a Poisson-Gaussian process fit of the nonlinear rate function, with its approximate 95% posterior band. The stimulus was projected onto the spike-triggered average (STA), and the projection and spikes were aggregated over the response intervals used by the Poisson count model. A random subset of observations is plotted as blue jittered dots to avoid overplotting. **(B)** Optimum nonlinear encoders for *L*_*p*_-norm reconstruction loss with mean firing rate constraint. Three of the four encoders use the same standard normal (Gaussian) prior: the thin grey curves at *p* = 2 and *p* = 7, drawn as solved under the mean rate constraint, and the thick curve at *p* = 5.75, the selected cell of that family. The remaining thick curve is the selected cell of the Student-*t* family (*ν* = 1.5) at *p* = 8.5, which matches the measured curve more closely. The two selected cells are compared to the measurement after fitting a gain and a baseline. **(C)** Predicted response marginals under the codes of panel B and the marginal distribution from spike data. The two selected cells predict nearly identical marginals, differing by at most 0.008 in probability, so the Student-*t* curve is drawn dashed over the Gaussian one; the response marginal does not discriminate between the two priors. See S1 Appendix, Numerical optimization and evaluation for the LNP model, for the optimization objective, the comparison criterion, and the prior-family analysis.

With a standard Gaussian prior matching the laboratory stimulus projections, we obtained the closest match to the empirical H1 curve at *p* = 5.75. However, this curve still overpredicted firing at large positive stimulus projections. We compared shapes after fitting gain and baseline, giving greater weight to log-rate discrepancies where the empirical Gaussian-process fit had lower posterior uncertainty. The remaining mismatch motivated us to ask whether a prior inspired by natural motion statistics would better describe the measured nonlinearity. In free-flying blowflies, the distribution of yaw velocities has a pronounced central peak and long tails, reflecting motion between saccades and during rapid turns [65].

We therefore considered a heavier-tailed Student-*t* prior, whose tail index *ν* controls the weight of the tails. The closest matches occurred at *p* = 8.5 for both *ν* = 1 and *ν* = 1.5, and both reduced the shape mismatch. The fits were nearly identical; we display *ν* = 1.5 because it is less affected by truncation of the numerical stimulus grid (Fig. 6B). The corresponding response distributions are neither uniform nor Poisson (Fig. 6C). The selected exponents describe agreement with this recording; they do not provide predictive estimates of the neural loss function or establish adaptation to a heavy-tailed natural ensemble.

This dataset was also analyzed by Wang and coworkers [57], who used a population-level tuning-curve framework with a Fisher-information approximation to the *L*_*p*_-norm loss. Our analysis differs in that we work with the exact posterior distribution under a single-neuron input-output nonlinearity.

## Discussion

We have synthesized Barlow’s efficient coding hypothesis with the Bayesian brain hypothesis in order to formulate a Bayesian theory of efficient neural coding. In this theory, an efficient neural code corresponds to an encoding model that produces optimal posteriors over stimuli under a capacity constraint on the neural response. The optimality of the posterior is determined by a loss function *L*, and the choice of loss function can have major effects on the optimal code. We have shown that Barlow’s original theory corresponds to a special case of this theory in which the loss function is the Shannon entropy of the posterior, corresponding to codes that maximize mutual information between stimulus and response. However, such codes may be inefficient with respect to loss functions that are sensitive to the shape of the posterior, or the size of different decoding errors.

To illustrate our framework, we have derived Bayesian efficient codes for two canonical neural coding problems: (1) population encoding of high-dimensional stimuli with linear receptive fields; and (2) single-neuron encoding of low-dimensional stimuli with a nonlinear response function. In the first case, we showed that the “whitening” solution favored by classic efficient coding can be sub-optimal for other loss functions, and that even in the high-SNR (signal-to-noise ratio) regime, optimal codes may *increase* the correlations between neurons. In the second case, we re-examined the classical results of Laughlin, and showed that the nonlinear response functions of the blowfly LMC neurons are in fact more consistent with a Bayesian efficient code that minimizes the average square root of the decoding error than a code that maximizes mutual information. In a complementary analysis of spiking responses from a blowfly H1 neuron under a linear-nonlinear-Poisson encoding model, we found that the measured nonlinearity is better matched by a finite-*p* code than by the infomax solution, but that the selected exponent depends on the stimulus prior and the curve-comparison criterion. While these examples are all highly simplified, they illustrate the power of this general formulation of efficient coding, and show surprising and non-intuitive results that contrast with previous findings.

The H1 result is therefore a phenomenological fit rather than an ecological account of the neural loss function. Relative to a Gaussian prior with squared-error loss, a heavier-tailed prior makes large stimulus projections more common, while a larger value of *p* makes large reconstruction errors more costly. Together, these changes shift coding resources toward the tails. One possible biological interpretation is that H1 is adapted to rare, abrupt optic-flow events for which large errors have disproportionate behavioral costs.

### Relationship to previous work

In the years since the seminal papers of Attneave and Barlow, a large literature has taken up the problem of optimal neural coding, spanning a wide range of both information and non-information-theoretic frameworks. Barlow’s original paper considered noiseless encoding, where each stimulus maps to a deterministic response [2]. Atick and Redlich extended this framework to incorporate noise, deriving the remarkable result that optimal receptive field shapes change with SNR, consistent with observed changes in retinal ganglion receptive fields [10]. They showed that whitening is optimal only at high SNR, and that optimal responses become correlated at lower SNR—note that this differs from our result showing that whitening can be sub-optimal even at high SNR for other loss functions (Figs. 3 & 4).

Several alternate versions of efficient coding based on information theory were advanced in the early years. Barlow first proposed that neurons optimize a quantity he called *redundancy*, given by 1 − *I*(**x, y**)*/C*, or one minus the mutual information divided by the channel capacity *C* [2]. (In fact, Barlow referred to his theory as the “redundancy reduction hypothesis”, and the more common “efficient coding hypothesis” label has only appeared more recently, e.g., [66]) Atick and Redlich modified Barlow’s theory to replace the *C* in the denominator by *C*_*out*_, a modified notion of channel capacity related to the system’s total power, which they held to be more biologically plausible [10, 40]. Neither theory therefore amounted to a pure “infomax” hypothesis, such as that advanced by [8], since optimality could be increased either by increasing mutual information or decreasing the channel capacity *C* or *C*_*out*_ in the denominator.

It bears mentioning that Barlow’s definition of redundancy has no relationship to the concepts of redundancy and synergy later introduced by Brenner and colleagues, which quantify whether groups of neurons encode more or less information jointly than separately [67–70]. An efficient code according to Barlow or Atick & Redlich may be perfectly efficient in the sense of maximizing the ratio of information to capacity, while being either redundant or synergistic in the (more widely used) sense defined by Brenner *et al* [71].

Theories of efficient coding based on information-theory have been applied to a wide variety of different sensory systems and brain areas. These can be roughly grouped into those focused on linear receptive fields [35, 38, 72, 73], those focused on tuning curves [25–28, 54, 74–77], and those addressing some aspect of population coding, such coupling strengths between neurons [78, 79]. A substantial portion of this literature work has approached the problem of optimal coding through the lens of Fisher information [25, 26, 28, 57, 74, 76], although Fisher information may not accurately quantify coding performance in low SNR settings (e.g., short time windows or low firing rates) [17, 24, 27, 80].

A substantial literature has also considered optimal coding under alternate loss functions, and recent work has shown that codes optimized for mutual information may perform poorly for non-information-theoretic loss functions [81]. The most well-studied alternate loss function is the squared loss, 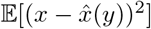, which results in so-called “minimum mean squared error” codes; such codes achieve minimum expected posterior variance (as opposed to minimum posterior entropy). These codes have received substantial attention in both engineering [82, 83] and neuroscience [15, 16, 18, 24, 27, 84, 85]. Optimal codes for a wide variety of other losses or optimality criteria have also been proposed, including: minimization of motor errors [14], maximization of accuracy [86–88], optimal coding for control applications [89], and optimal future prediction [12, 90–93], and loss based on natural selection [6]. Other recent work has considered codes that maximize metabolic efficiency [11, 13, 20, 94, 95], which in our framework corresponds more naturally to the constraint (which is concerned with use of finite resources) than the loss function (which is concerned with the posterior distribution).

Since the original preprint version of this manuscript was posted in 2017, the Bayesian efficient coding framework has been taken up across a range of problems in sensory coding, decision-making, and cognition. Representative examples include applications to the efficient coding of subjective value [96], visual working memory [97], psychoeconomic judgment [98], population coding with nonlinear mixed selectivity [99], functional diversity among sensory neurons [75], adaptive coding for dynamic sensory inference [19], the statistical analysis of neural-system optimality [100], and as a reference framework in reviews of adaptive neural coding [101]. We take this uptake as evidence that the four-ingredient decomposition articulated here – prior, encoding model, capacity constraint, and loss functional – captures a useful set of abstractions for reasoning about normatively optimal neural codes, beyond the specific examples we develop below.

In the statistics and decision-making literature, our work is closely related to Bayesian statistical decision theory [102–104], although such work has tended not to consider the problem of optimal sensory coding. One noteworthy difference between our framework and classical Bayesian decision-making theory is that the loss functions in decision theory are typically defined in terms expected cost of a point estimate (e.g., mean squared error or percent correct). In our framework, by contrast, the loss is defined as a *functional* of the posterior, that is, an arbitrary integral over the posterior distribution. This allows us to consider a wider class of loss functions, such as the posterior entropy or the average posterior standard deviation, which cannot be written in terms of an expected loss for a point estimate of the stimulus. Thus, our loss function merely specifies what counts as a good or desirable posterior distribution, without assuming how the posterior will be used (e.g., to generate a point estimate). This distinction also separates our framework from more recent unifying treatments of efficient coding and Bayesian inference that remain within a point-estimate decision-theoretic view, such as the *L*_*p*_-based frameworks of [57] and [29].

### Relevance of Shannon information theory

Shannon’s information theory holds a special status in engineering, signal processing, and other fields due to its universal implications for communication, compression, and coding in settings of perfect signal recovery. However, it is unclear whether information-theoretic quantities like entropy and mutual information are necessarily relevant to information processing and transmission in biological systems [105]. In particular, Shannon’s theory (1) requires complex computation and long delays to encode and decode in ways that achieve optimality [5]; (2) ignores biological constraints (e.g., neurons cannot implement arbitrary nonlinear functions); (3) applies to settings of perfect signal recovery, which may not be possible or even desirable in biological settings.

In the Bayesian efficient coding, the encoder is selected from a biologically relevant parametric family. And although we consider the (negative) mutual information as one possible choice of loss function, there is no *a priori* reason for preferring it to other loss functions; as we have shown, optimizing MI will not necessarily give good performance for other losses (e.g., Figs. 2 & 3). We therefore find no justification for claims (commonly found in the neuroscience literature) that, in the absence of knowledge about the brain’s true loss function, it is somehow best to maximize mutual information.

A framework more relevant to sensory coding in the nervous system is Shannon’s rate distortion theory for lossy encoding [31]. Given an allowed maximum distortion (as a measure of encoding-decoding error) *L*^∗^, the rate distortion function *R*(*L*^∗^) describes the minimum necessary mutual information of the channel (implemented by the encoder-decoder pair) to achieve it: 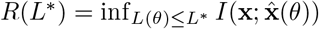. One can prove that the more mutual information is allowed, the smaller the achievable distortion; in fact *R*(*L*^∗^) is a non-increasing convex function [32]. This provides an interesting overlap between the two theories: a subclass of BEC with mutual information constraint can be reformulated as a special case of rate distortion theory. Consider the following dual formulations:

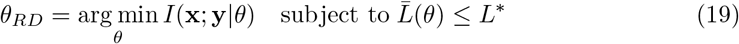

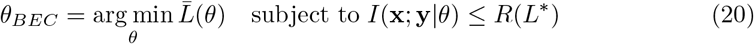

where 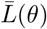 is the loss given in eq. 1. The rate distortion solution *θ*_*RD*_ coincides with the BEC solution *θ*_*BEC*_, because the distortion function is monotonically decreasing. Although this provides a insightful connection, such relations do not always exist, and more generally it is not clear why a bound on mutual information would be a biologically relevant constraint on a sensory encoder.

### Application to experimental data

In this manuscript, we used the BEC framework to identify loss functions consistent with neurophysiology datasets. To accomplish this, we searched for the optimal encoding model under different choices of loss function, and asked which of the resulting encoding models most closely resembled the empirical encoding model. Although both of our examples came from the early visual system, one could apply the same approach to datasets from any sensory modality (e.g., audition, olfaction, somatosensation). This would reveal what loss function (if any) the neurons in different brain and different modalities seek to optimize. Researchers can therefore test whether neural representations maximize mutual information (as posited by classic efficient coding) or some other loss metric.

A second way to apply the BEC framework to data is to identify the prior distribution for which an observed sensory encoding is optimal (cf. [47, 106, 107]). In this case, we would take the encoding model parameters inferred from neurophysiological data and optimize the prior distribution under different fixed choices of loss functions or constraints. The inferred priors may then serve as predictions of the distribution of natural signals for which the system is optimized, or as an assay of neural adaptation in experiments with time-varying stimulus statistics.

Finally, the BEC framework could be used to test which constraints are most relevant to a particular neural code. Here we assumed that the relevant constraint was on the range (for analog responses; Fig. 5), or on the mean firing rate (spiking responses; Fig. 6). However, one could test whether observed responses are more consistent with alternate sets of constraints, including metabolic or energetic constraints [13, 106, 108], constraints on higher-order response statistics [109], or architectural constraints on the model parameters (e.g. sparsity, wiring length) [78, 110, 111].

The general domain of application of BEC is thus the same as classic efficient coding; the key difference is that BEC has one additional “knob” that can be tuned, namely the loss function. As classical efficient coding provided new explanations for adaptation and sparse coding, the BEC framework can be used to extend and refine such hypotheses, set free from the restrictive assumption that mutual information is the relevant loss function for all of behavior.

### Limitations and future directions

What cost or loss function is the nervous system optimizing for? We emphasize that BEC alone cannot serve as a normative theory to answer this question; each problem and environment should dictate the loss functional and prior. Rather, BEC should be used as a guiding principle that frees the theory of efficient coding from its traditional reliance on information theoretic principles, and to cover an appropriately broad range of theories of optimal neural encoding.

BEC on its own has many degrees of freedom and is likely under-constrained by current neural and behavioral observations. Moreover, different combinations of constraints and loss functions may be consistent with a single encoding model, providing multiple “optimal” explanations for a single encoder. The priors, encoding models, constraints, and loss functions we have considered here were guided primarily by tractability as opposed to neural or biological plausibility, but some of these components (e.g., prior and noise) can be quantified with measurements [41, 112, 113]. We can start to make reasonable inferences about constraints and loss functions through careful study of the brain’s resource consumption [114, 115] or the behavioral consequences of different kinds of errors [116–118].

A variety of other issues relating to normatively optimal neural coding remain to be addressed by our theory:

#### Computational demands

the BEC framework does not consider constraints on the brain’s computational capabilities. In particular, our theory—like with the Bayesian brain hypothesis itself—assumes that the brain can compute the desired posterior distribution over stimuli given the response, or at least extract the posterior statistics needed for optimal behavior. This is almost certainly not the case for many of the complex inference problems the brain faces. It seems more likely that the brain relies on shortcuts, heuristics, and approximate inference methods that result in sub-optimal use of the stimulus information contained in neural responses relative to a Bayesian ideal observer [119]. The BEC paradigm could therefore be extended to incorporate computational constraints: for example, restricting the readout of neural activity to be linear, or the use of particular approximate Bayesian inference methods [120].

The BEC framework does not require a particular neural representation of the posterior. For example, a downstream population could use a probabilistic population code [22], a sampling-based representation [121], or a distributed distributional code that represents selected conditional expectations rather than an explicit probability density [122]. If this population preserves the posterior quantities required by the loss, introducing it does not alter the BEC optimum; it provides a neural implementation of the readout already assumed by the theory. If its representation is noisy, capacity-limited, or approximate, however, those constraints must enter the optimization and can change the optimal upstream code.

#### The role of time

we have focused our analyses on coding problems that follow the sequential nature of Bayesian inference (stimulus → response → posterior distribution) and have ignored the continuous-time nature of stimuli, responses, and behavior. However, there is nothing about our theory that precludes application to continuous-time problems, and previous work has formulated population codes capable of updating a posterior distribution in continuous time [123–127]. In many cases optimal coding depends on the timescale over which information is integrated: for example, previous work has shown that optimal tuning curve width depends on the time over which spikes are generated, with longer time windows necessitating narrower tuning [17, 27]. Recent work has discovered limits on fast timescale transmission of information in physical and biological systems [128], but there is still a need for a theory of optimal coding in settings where there is uncertainty about the time (or times) at which the posterior distribution will be evaluated or “read out”.

#### Latent variables

we have not so far discussed latent variable models, in which there are additional unobserved stochastic components affecting either the stimulus or the neural response. In stimulus latent variables models, the quantity of interest is a latent variable **z** that may be regarded as underlying or generating the sensory stimulus **x**. (For example, **z** might the velocity of an object or the identity of a face, and **x** is the resulting image or image sequence presented to the retina [129]). In such settings, the posterior over the latent variable **z** might be the quantity of interest for an optimal code; BEC can handle this case by defining a loss function sensitive to the posterior *p*(**z**|**y**). This can be seen as a valid instance of BEC because this distribution can be written as a functional of the stimulus posterior: *p*(**z**|**y**) = ∫ *p*(**z**|**x**)*p*(**x**|**y**) d**x**, where *p*(**z**|**x**) is the posterior over the latent given the stimulus under the generative model. Latent variables also arise in models of neural activity with shared underlying variability [130–134]. In such cases, it is natural to write the encoding model itself as a joint distribution over activity and latents, *p*(**y, z**|**x**); the BEC paradigm can once again handle this case by marginalizing over the latent to obtain *p*(**x**|**y**). Thus, BEC can accommodate both kinds of latent variable models typically used in neuroscience, and the consideration of stimulus-related latent variables may motivate the design of loss functions that are sensitive to particular latent variables while discarding other aspects of the stimulus (e.g., as in the information bottleneck [92] or accuracy maximization analysis [86, 129]).

#### Transformation dependence

Shannon’s differential entropy, *L*_*p*_ reconstruction loss, and covtropy are *not* invariant to transformations of the stimulus variable. That is, it is important to measure what is meaningful in the right scale. For example, the LMC analysis was done in the Weber contrast. The results would have been different if it was transformed to a linear scale.

Although we have focused on two canonical problems that arise repeatedly in the neural coding literature, namely optimal linear receptive fields and optimal nonlinear input-output functions, we hope that future work will address other coding problems relevant to information processing in the nervous system (e.g., multi-layer cascades, dynamics, correlations), and will be extended to non-Bayesian frameworks (e.g., that take into account computational costs or constraints). We believe that the BEC paradigm provides a rigorous theoretical framework for neuroscientists to evaluate neural systems, synthesizing the Bayesian brain and efficient coding hypothesis into a formalism that can incorporate diverse optimality criteria beyond classic information-theoretic costs.

## Methods

### Bivariate Gaussian encoding example (Fig. 2)

The posteriors obtained in our first toy example rely on the well-known relationship between prior and posterior variance in a linear Gaussian encoder: 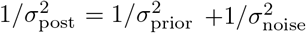. To achieve the posteriors shown in Fig. 2, we assumed following variances for the independent encoder noise in each dimension: **Encoder 1**: 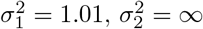; **Encoder 2**: 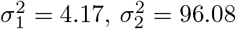; **Encoder 3**: 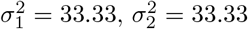.

### Blowfly LMC data from [30]

We extracted the data points from the figures in the original paper [30] to fit the models in Nonlinear response functions and for the plots in Fig. 5. Note that Laughlin’s stimulus variable is Weber contrast, 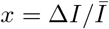: the deviation of a sample from the mean intensity 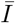 of the scan interval containing it, divided by that mean [30]. Therefore, the contrast variable is bounded below by −1 and unbounded above, so that the support of the contrast prior is [−1, ∞).

To fit the stimulus CDF and the response nonlinearity of the blowfly LMC, we fit a 3-parameter Naka-Rushton function |(*x* − *a*)^*b*^|*/*(*x* − *a*|^*b*^ + *c*) (plotted as grey dotted line in Fig. 5B) to estimate the stimulus CDF (Fig. 5A). The parameters for the stimulus CDF were *a* = −1.52, *b* = 5.80, *c* = 7.55, and *a* = −1.90, *b* = 5.84, *c* = 37.17 for the LMC response.

### Closed-form optimal nonlinearity for the LMC prior

The Naka-Rushton CDF fit above is a log-logistic prior, for which the numerically optimized nonlinearity has a closed-form counterpart in the limit of many response levels. Writing *F* for the fitted prior CDF and *b* for the fitted Naka-Rushton exponent, the *L*_*p*_-optimal nonlinearity is a beta CDF warp of *F*,

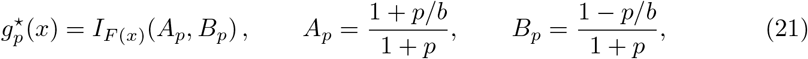

where *I*_*u*_(*A, B*) is the regularized incomplete beta function, that is, the CDF of Beta(*A, B*) evaluated at *u*. The exponent enters only through the two numbers *A*_*p*_ and *B*_*p*_. At *p* = 0 they are both 1, the warp is the identity, and 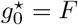 recovers histogram equalization, so infomax is the *p* → 0 member of the family. A solution exists only for 0 ≤ *p < b*, which is exactly the range over which the prior has a finite *p*-th moment. The derivation, the affine rescaling induced by truncating the fit at the contrast floor (the fit has *a <* −1), and a numerical check of eq. 21 against the finite-level quantizer optima are given in S1 Appendix, Closed-form optimal nonlinearity for a log-logistic prior.

### Blowfly H1 data from [63]

We projected the stimulus onto its spike-triggered average (STA). Spikes were summed over sliding 50 ms windows, and the corresponding stimulus projections were averaged over the same windows. We estimated the empirical nonlinearity (Fig. 6A,B) using Gaussian-process-Poisson regression [64]. The conditional spike count was Poisson with an exponential link, and the latent log rate had a Gaussian-process prior with a linear mean and an isotropic squared-exponential covariance function. We used a Laplace approximation to obtain the posterior mean and variance of the log rate. The posterior mean defines the measured nonlinearity, while the posterior variance defines its uncertainty band and weights the shape comparison below. The candidate nonlinearities were monotone and non-negative over the range of stimulus projections supported by the data. Similarly with the LMC data, we then optimized the nonlinearity under different loss functionals with the marginal firing rate constraint (Fig. 6B). The optimization uses the exact finite-count posterior and constrains the mean count to its measured value. We compare each predicted nonlinearity with the measured one after fitting a gain and a spontaneous rate. These two parameters were fitted using non-overlapping windows. S1 Appendix, Numerical optimization and evaluation for the LNP model gives the objective, the shape comparison, and the sensitivity analysis over the assumed stimulus prior.

## Data availability statement

Data and replication code, in both MATLAB and Python, are available at Zenodo (DOI:10.5281/zenodo.3970841), which resolves to the most recent version of the record.

## Acknowledgments

We would like to thank Jacob Yates, Ozan Koyluoglu, and Kaining Zhang for helpful comments and discussions.

## Author contributions

**Conceptualization:** IMP, JWP. **Formal analysis:** IMP, JWP. **Investigation:** IMP, JWP. **Methodology:** IMP, JWP. **Software:** IMP. **Visualization:** IMP. **Writing – original draft:** IMP, JWP. **Writing – review & editing:** IMP, JWP.

## S1 Appendix. Supporting information

### Loss functionals

Loss functionals quantify the goodness of the posterior distribution *P* (**x**|**y**, *θ*). There are broadly two classes of loss functionals: **entropic loss** or **reconstruction loss**. Entropic losses quantify the average uncertainty of the posterior: The more concentrated the posterior, the better.

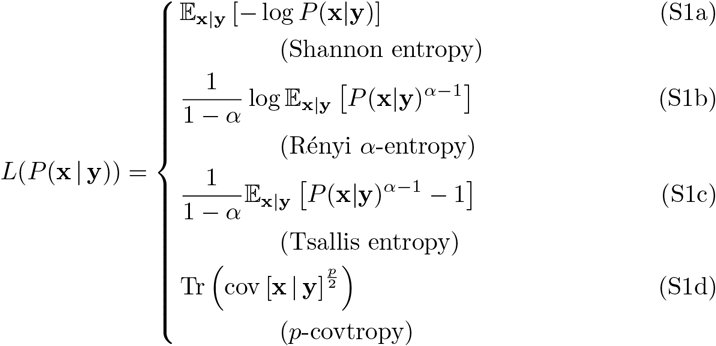

Both eq. S1b and eq. S1c converge to eq. S1a in the limit of *α* → 1.

Traditional entropic losses do not discriminate points in the stimulus domain, that is, any error is treated equally. However, when stimulus space is ℝ^*n*^, it makes sense to prefer locally tight posterior distributions. One such measure is the *p*-covtropy which measures the posterior concentration around the posterior mean (only defined for distributions in ℝ^*d*^). Note that 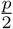 ‘th power of the posterior covariance matrix corresponds to taking *p*’th power of the standard deviations along each principal axis through diagonalization.

The second class of loss functionals depends on a specific readout (a.k.a. estimator or reconstruction) 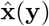 that maps the neural response **y** back to the stimulus domain. We consider a general average reconstruction error loss functional of the form: 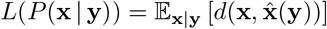 where *d*(·, ·) is a distortion measure.

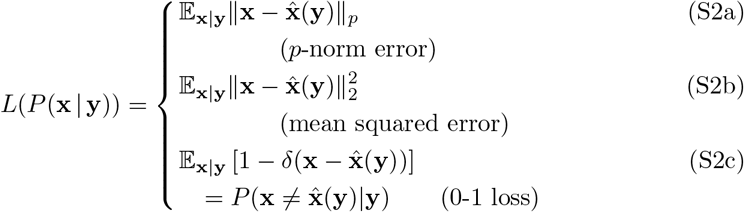

Mean squared error and the 0-1 loss are widely used in communication theory, machine learning, and statistics in general.

There are loose bounds that connect the posterior entropy with Bayesian error rate (e.g. Fano bound for classification) [83, 135, 136]. However, these bounds do not justify infomax strategy because better Bayesian error rate can be achieved with a system that directly optimizes for the target error measure.

Note the similarity between *p*-norm error and the *p*-covtropy when posterior mean is used for decoding. They coincide when *p* = 2, however for *p* ≠ 2, unlike the *p*-norm error, *p*-covtropy is invariant under unitary transformation of the stimulus space (which is the key difference between the 2D Gaussian and the linear receptive field examples). This is an important distinction if axes in **x** do not have special meaning, and rotated posteriors are considered equally good. We can make this connection rigorous for a Gaussian posterior, **x**|**y** ~ *N* (*µ, C*). Let *C* = *UDU*^−1^ be the eigendecomposition of the covariance matrix. Let *Z* ~ *N* (0, *D*) be aligned on the principal axes, the *p*-th power of the *p*-norm is given by,

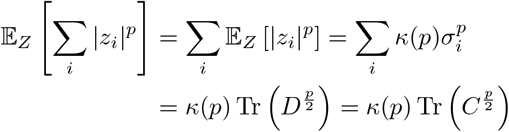

where 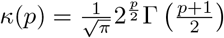.

In Linear receptive fields, we discuss the loss functions only for Gaussian posteriors. However, covtropy is well defined for any distribution with a valid covariance matrix. Note that minimizing the limiting case of covtropy for *p* → 0 and maximizing mutual information do not coincide in general since posterior entropy is not a sole function of the covariance in general.

### Equivalence of MSE and covtropy for *p* = 2

Here we provide a simple proof showing that minimizing covtropy for *p* = 2 corresponds to minimizing mean squared error (MSE). Let 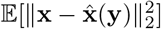 denote the MSE for any estimator 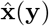, where expectation is taken with respect to the posterior *P* (**x**|**y**). It is well known that the posterior mean or “Bayes least squares” estimator 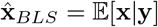 achieves the minimum of the MSE, which is then given by 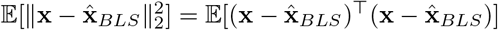. We can use identities involving trace and the definition of covariance to show that MSE is equal to 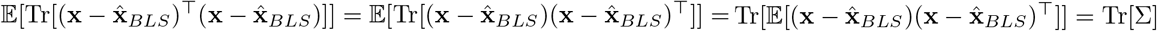, which is the covtropy with *p* = 2. Thus, the code that achieves minimum *p* = 2 covtropy corresponds to the code that achieves minimum MSE point estimation of the stimulus.

### Linear receptive fields

Here we derive the optimal receptive fields for linear encoding under Gaussian noise (see Linear receptive fields). Recall the model (eqs. 3-5):

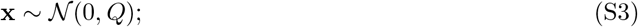

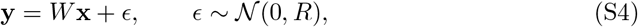

and we wish to find the optimal *W* subject to the power constraint

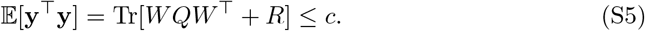

We make the assumption that *Q,R*, and *W* matrices commute, which means they share a common eigenbasis or can be diagonalized by the same orthogonal matrix. This occurs, for example, if all three are circulant matrices, so that *W* consists of shifted copies of a single RF shape.

For Fig. 4, *Q* is the 100-dimensional projection of a 1*/f* ^2^ stimulus spectrum and *R* = 0.001*I*, giving irreducible noise power Tr(*R*) = 0.1. The high-, intermediate-, and low-power conditions use *c* = 0.655Tr(*Q*) = 99.5643, 0.5, and 0.2, respectively. The all-active closed form in the main text applies while the matrix inside its square root is positive semidefinite. At lower power, the constrained optimum sets some gains to zero and reallocates power across the remaining modes. At intermediate power, the infomax and *p* = 1, 2, 4 encoders retain 100, 100, 100, and 57 active modes; at low power they retain 100, 76, 38, and 23.

#### Infomax

Maximizing information is equivalent to minimizing the posterior entropy:

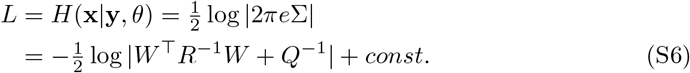

We can find the optimal *W* subject to the power constraint above using the method of Lagrange multipliers:

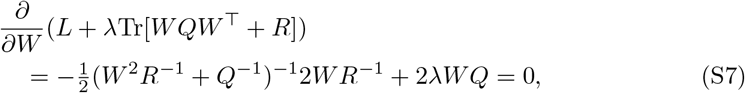

where *λ* is a Lagrange multiplier. This implies

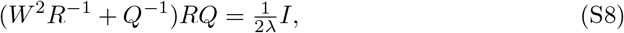

giving

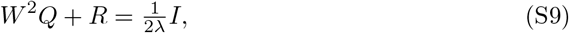

and finally

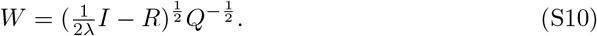

Plugging this solution into the constraint (eq. S5) gives

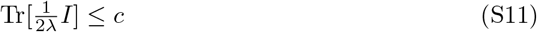

which implies 1*/*(2*λ*) = *c/n*, where *n* = dim(**y**) is the number of neurons. Substituting for *λ* gives the desired expression:

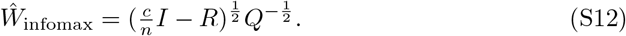

#### Minimum p-Covtropy

We can take a similar approach for the loss function we have called *p*-covtropy, which is effectively the mean *p*’th power error in the eigenbasis of the posterior distribution. This is given by

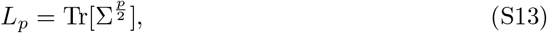

which involves the posterior covariance matrix Σ to the matrix-power *p/*2. It turns out this loss function has the same optimum as infomax loss in the limit *p* → 0.

Once again, the method of Lagrange multipliers allows us to solve for *W* explicitly in the case that *W, R*, and *Q* commute. Taking the derivative of the Lagrangian with respect to *W* and setting to zero yields

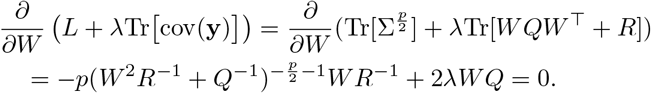

This can be simplified to

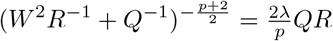

and so

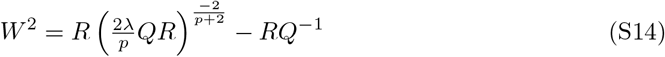

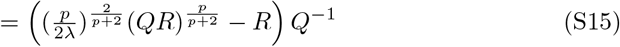

and finally

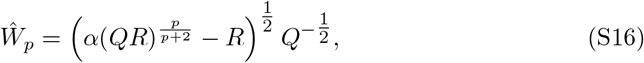

with 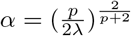. Substituting *Ŵ*_*p*_ into the power constraint (eq. S5) gives:

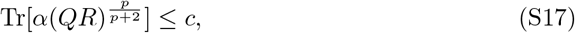

which achieves the constraint with equality when

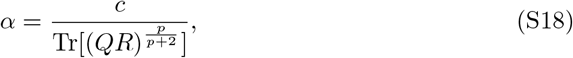

which completes the derivation.

### Closed-form optimal nonlinearity for a log-logistic prior

When the stimulus prior is log-logistic, which the Naka-Rushton family fit to Laughlin’s contrast distribution is (Methods), the optimum can instead be written down in closed form in the limit of many response levels. The exponent *p* then enters only through the two parameters of a beta distribution, and a solution exists only for *p* below the tail index of the prior.

Let *g* be an increasing nonlinearity of the Weber contrast *x* ≥ −1, mapping it onto the response range normalized to [0, 1], so that *g*(− 1) = 0 and *g*(∞) = 1. Its derivative *ρ* = *g*^*′*^ is a probability density on the stimulus axis: *ρ*(*x*) is the fraction of the response range spent per unit stimulus near *x*, so it sets the local resolution of the encoder, and ∫ *ρ* = 1 is the range constraint itself.

In the high-resolution limit, where precision is limited by a large but finite number of response levels, the local cell width is proportional to 1*/ρ*(*x*), and the average reconstruction error (eq. 16) takes the form *L*^*p*^ ∫ ∝ *P* (*x*) *ρ*(*x*)^−*p*^ d*x* to leading order. Minimizing it subject to ∫ *ρ* = 1 gives the classical optimal point density of high-resolution quantization [137–139], with the asymptotic theory for a prior on an unbounded support due to Zador [140]:

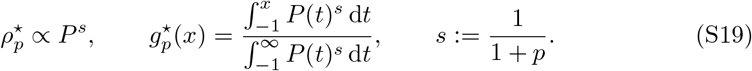

The case *p* = 2, *ρ*^⋆^ ∝ *P* ^1*/*3^, is the Panter-Dite rule [138]. Raising *p* flattens the resolution profile, moving resolution away from the mode of the prior and into the tails, where errors are large.

It is convenient to describe a nonlinearity by how it warps the uniformized stimulus *u* = *F* (*x*), which is uniform on [0, 1], where *F* is the prior CDF. Every increasing *g* with *g*(− 1) = 0 and *g*(∞) = 1 can be written as *g* = *W* ◦*F* for a single increasing warp *W* : [0, 1] → [0, 1] with *W* (0) = 0 and *W* (1) = −1. The Naka-Rushton CDF fit to the contrast distribution (Methods), with its location at the contrast floor, *a* = −1, is exactly a log-logistic prior: log(1 + *x*) then has a logistic distribution with shape *b*, and in the uniformized coordinate the density is a product of powers of *u* and 1 − *u*,

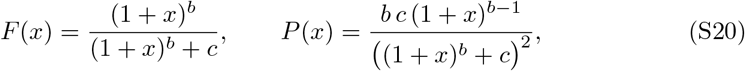

with *b* and *c* the fitted Naka-Rushton exponent and semisaturation, and

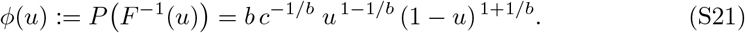

Under the change of variables *u* = *F* (*x*), so that d*u* = *P* (*x*) d*x*,

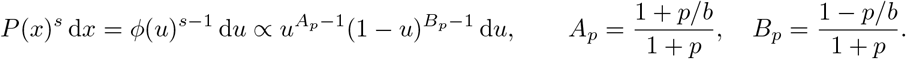

Thus, the integrals in (eq. S19) are incomplete beta integrals. The optimal warp is therefore the CDF of a beta distribution,

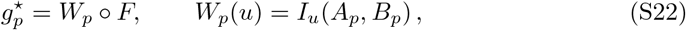

where *I*_*u*_(*A, B*) is the regularized incomplete beta function, that is, the CDF of Beta(*A, B*) evaluated at *u*. The entire dependence on *p* is carried by the two numbers *A*_*p*_ and *B*_*p*_ (Fig. S1A,B).

The same two numbers determine the response distribution. The response is *y* = *W*_*p*_(*u*) with *u* uniform, so its density is 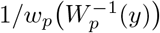, where *w*_*p*_ is the Beta(*A*_*p*_, *B*_*p*_) density. Here *A*_*p*_ *<* 1 exactly when *b >* 1, that is, when the prior has an interior mode, while 0 *< B*_*p*_ *<* 1 for every 0 *< p < b*; with both parameters below 1 the beta density is U-shaped, so *P* (*y*) is peaked at intermediate response levels (Fig. S1C, and Fig. 5C). Its peak sits where *w*_*p*_ is smallest, at *u* = (*b* − 1)*/*(2*b*), which is the prior CDF evaluated at the mode of the prior. For every *p >* 0, the most frequently used responses therefore encode the most probable stimulus.

They also give both ends of the family. Since *A*_0_ = *B*_0_ = 1 and Beta(1, 1) is uniform, *W*_0_ is the identity and 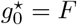, which is histogram equalization: infomax is the *p* → 0 member of this family rather than a separate criterion [30, 56]. At the other end, beta normalization requires *A*_*p*_ *>* 0 and *B*_*p*_ *>* 0. The first holds for every *p >* 0, whereas the second holds only for *p < b*. This is also the moment condition on the prior: since *P* (*x*) falls as *x*^−(*b*+1)^, E|*x*|^*p*^ *<* ∞ exactly when *p < b*. Thus, *no bounded-range code has finite L*_*p*_ *error for p* ≥ *b*. As *p* approaches *b* from below, the optimum spends an ever larger share of its response range on stimuli beyond any fixed contrast (Fig. S1).

The Naka-Rushton fit quoted in Methods has a free location *a* ≤ −1 rather than *a* = −1, and the physical contrast floor then truncates it. Writing *F*_0_ for the untruncated fit, which is log-logistic in *x* − *a* with the same shape *b*, the prior is

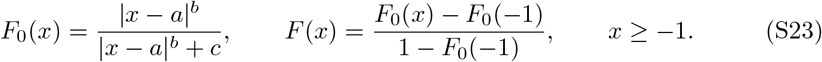

Truncation multiplies *P* by a constant on the support, which cancels in (eq. S19), so *A*_*p*_ and *B*_*p*_ are unchanged and the only difference is an affine rescaling of the same warp,

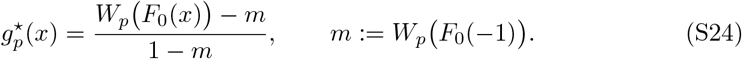

The case *a* = −1 has *F*_0_(− 1) = 0 and *m* = 0, which returns eq. S22. Fig. S1 shows this family for the fitted prior.

The closed form above is the high-resolution (*N* → ∞) limit of the exact finite-*N* quantizer problem it was derived from. Fig. S2 checks it directly against that finite-*N* problem, solved independently by Lloyd-Max fixed-point iteration [141, 142] rather than by the formula being checked.

**Figure S1.**
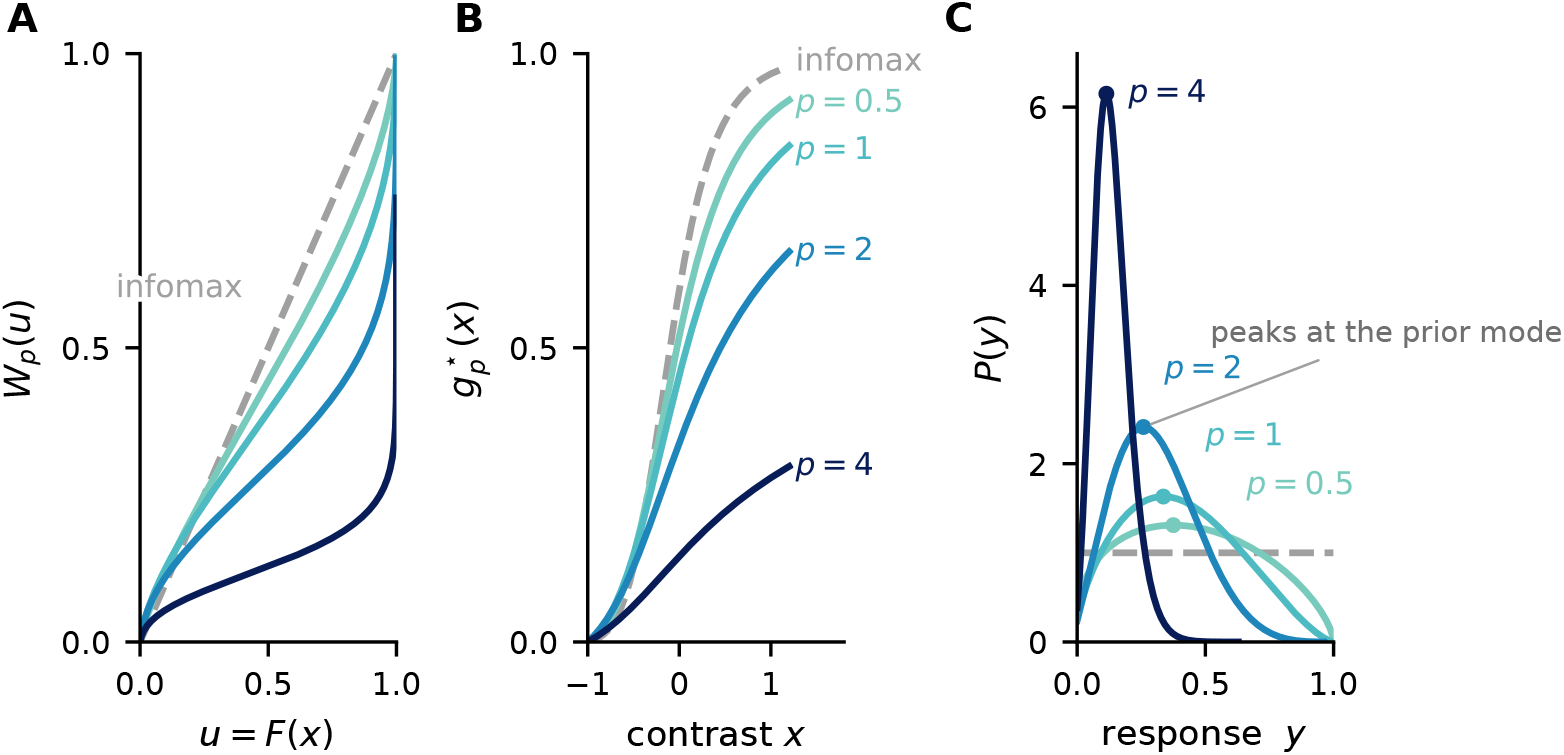
Increasing *p* shifts coding resolution from common contrasts toward the tails of the stimulus distribution. **(A)** Optimal warps *W*_*p*_ of the uniformized stimulus *u* = *F* (*x*) for the fitted contrast prior. The identity warp at *p* = 0 is the infomax solution; increasing *p* assigns more of the response range to rare, large contrasts. **(B)** The same solutions in contrast coordinates, 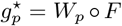. **(C)** Response distributions induced by each code. Infomax uses every response equally often, whereas each *p >* 0 solution is peaked at the response assigned to the most probable stimulus. The largest displayed exponent shows the limiting behavior as *p* approaches the tail index *b*, beyond which no finite-error solution exists.

### Numerical optimization and evaluation for the LNP model

The LNP optima of Fig. 6B have no closed form. Let *x* denote the STA-projected stimulus and *g*(*x*) the conditional mean spike count. For each count *y*, we use the exact Poisson posterior *P* (*x*|*y*) ∝ Poisson(*y*; *g*(*x*)) *P* (*x*) and its Bayes estimator under *L*_*p*_ loss.

We minimize the response-averaged reconstruction error,

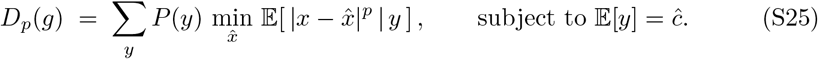

Here *ĉ* is the empirical mean count. We optimize over non-decreasing, non-negative *g* from multiple starts.

The mean-rate optimum has zero baseline, whereas H1 fires spontaneously. Before comparing shapes, we therefore fit the affine map *λ*(*x*) = *a g*(*x*) + *b* by Poisson maximum likelihood. Let 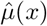 and 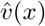 denote the posterior mean and variance of the log rate under the Gaussian-process-Poisson fit (Blowfly H1 data from [63]). We compare the transported model with this fit using

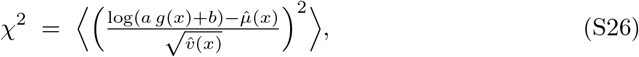

where the brackets denote an average over the data-constrained part of the measured curve.

The normalized STA projection *x* is approximately standard normal in this experiment, so a standard-Gaussian prior matches the laboratory stimulus. The best-fitting standard-Gaussian solution nevertheless overpredicts H1 at large positive projections. Natural-flight yaw velocities have a sharp central peak and long tails produced by intersaccadic and saccadic motion [65], and previous work modeled this same H1 recording with a power-law speed prior [57]. These results motivate, but do not determine, broader or heavier-tailed adaptation priors for the STA projection. We therefore vary Gaussian width and Student-*t* tail index (Fig. S4). The Student-*t* family was scaled to match the interquartile range of the standard normal, isolating tail shape from central scale. Across both families, *p* changes the predicted nonlinearity more than the prior parameter, and the best matches remain far from the infomax limit.

**Figure S2.**
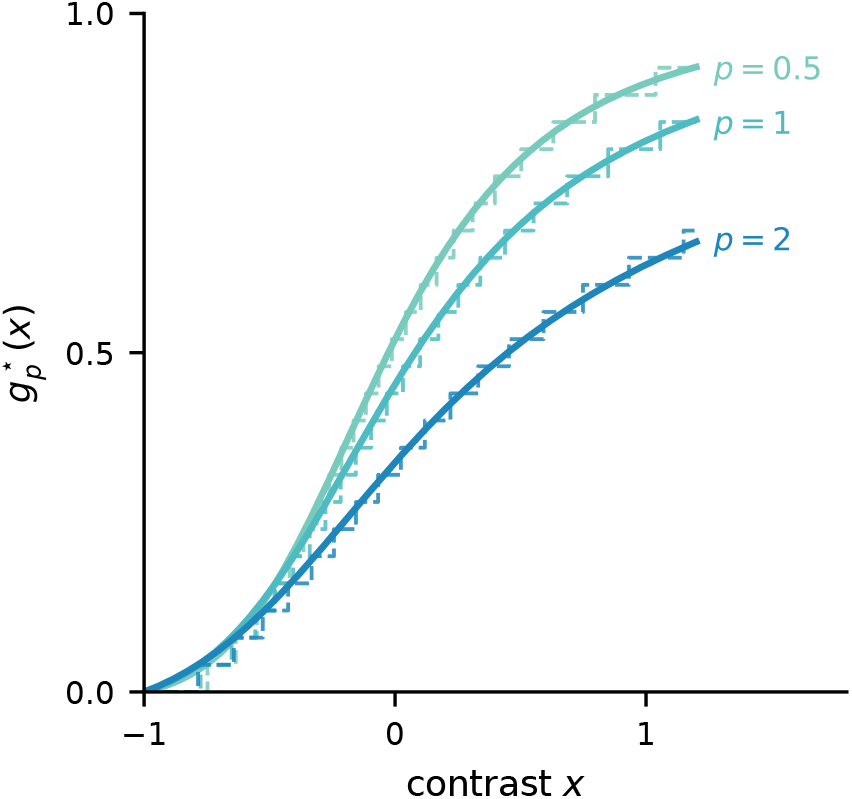
The closed-form nonlinearities agree with independently optimized finite quantizers. The high-resolution solution 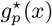 (solid) is compared with an independently optimized finite-level Lloyd-Max quantizer for the same prior and exponent (dashed staircase). Their close agreement shows that the closed form describes the optimal code well at finite response resolution.

A Student-*t* prior has no finite *p*-th moment for *p* ≥ *ν*, so those solutions depend on restricting the stimulus range to that supported by the data. We display *ν* = 1.5 because it is heavy-tailed but places less mass outside this range than the more extreme priors. The improved match under heavy tails therefore does not establish that H1 is adapted to a heavy-tailed natural ensemble.

**Figure S3.**
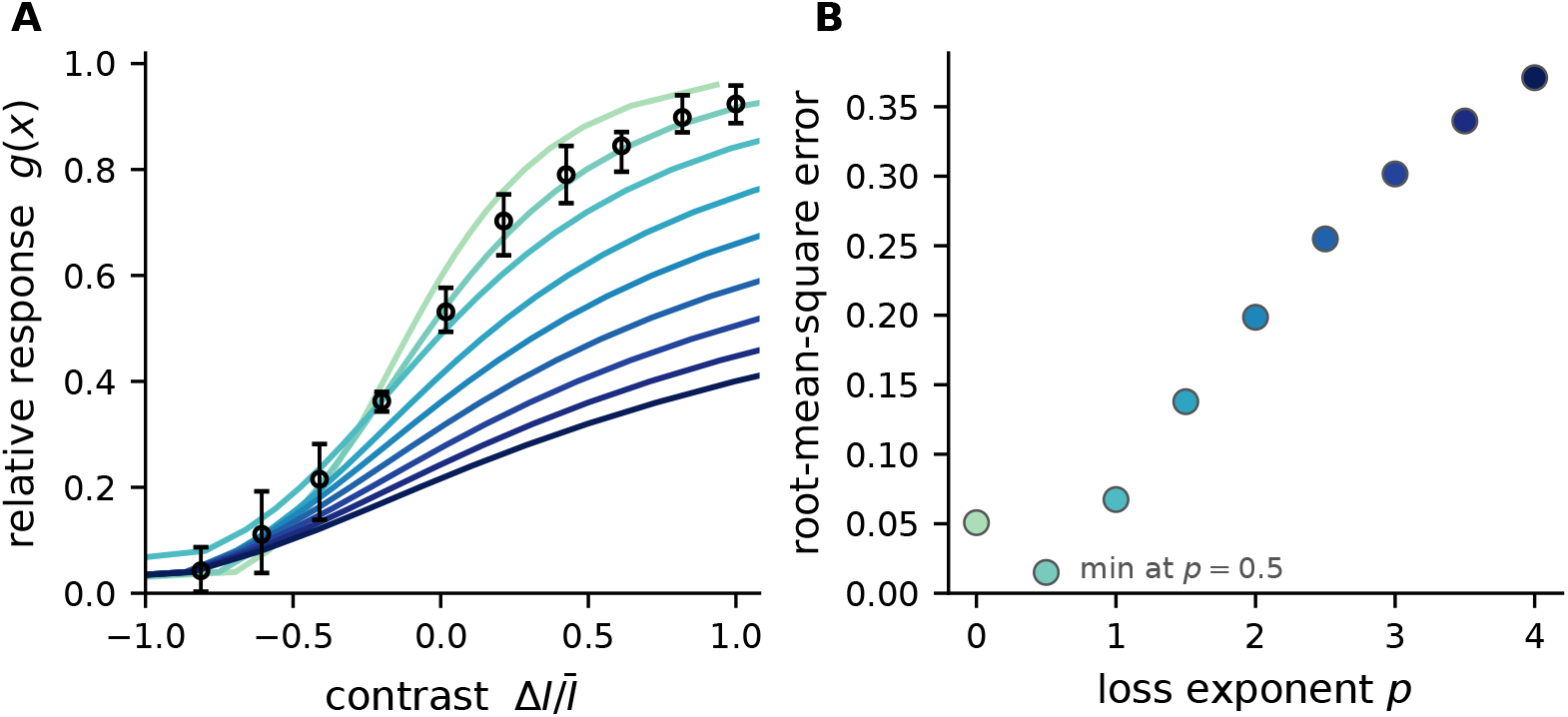
The LMC response favors 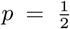 over infomax and larger loss exponents. **(A)** Optimal nonlinearities under *L*_*p*_ loss, computed from the fitted stimulus distribution and compared with the measured LMC response (circles, mean *±* error bars). The *p* = 0 curve is the infomax solution; color encodes *p*. **(B)** Root-mean-square error (RMSE) between each predicted nonlinearity and the measured response. The RMSE falls from 0.051 under infomax to 0.015 at 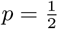, then increases monotonically for larger *p*.

**Figure S4.**
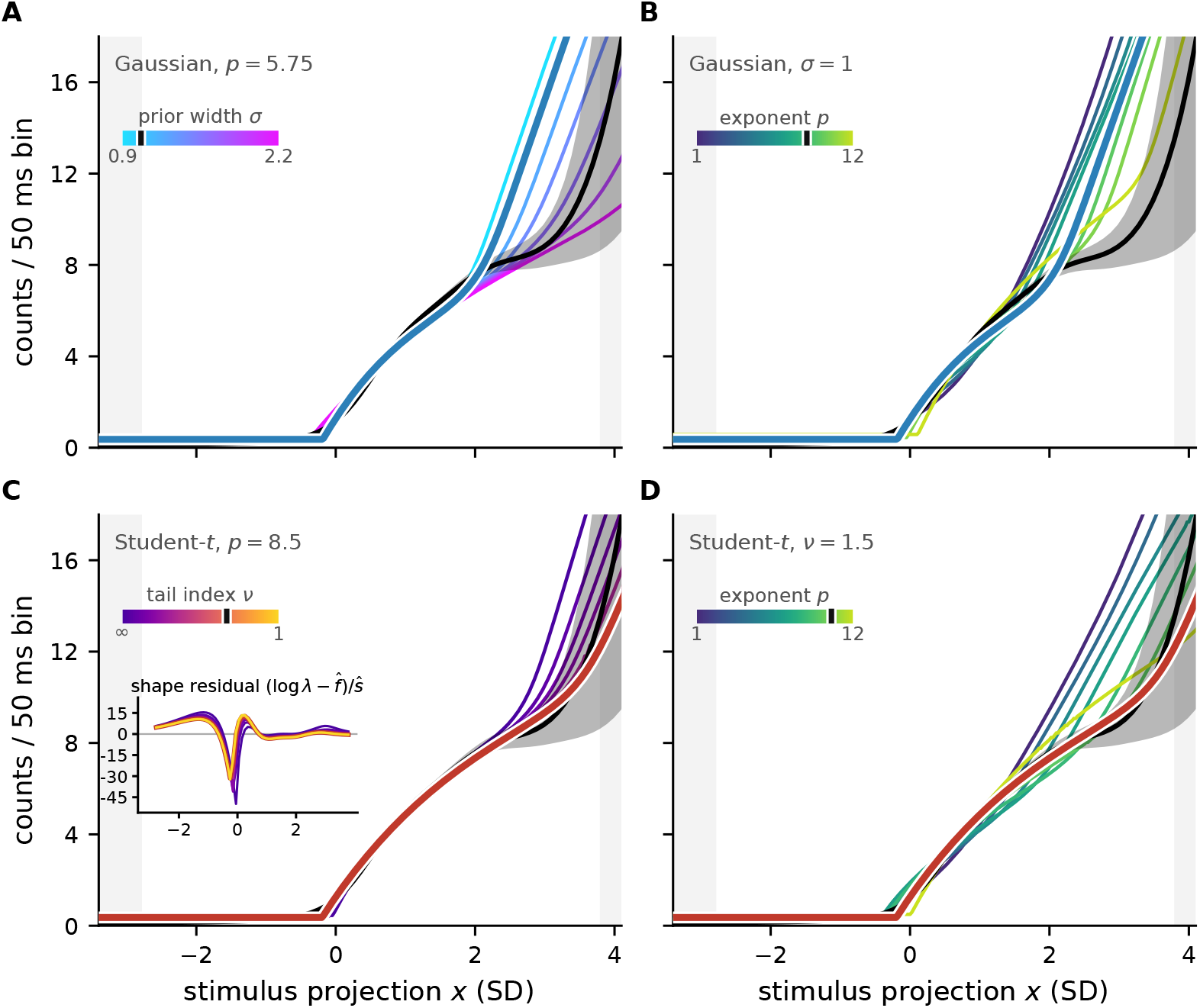
The H1 nonlinearity favors a large loss exponent across Gaussian and heavy-tailed stimulus priors. Optimal LNP nonlinearities (colored curves) are compared with the measured H1 nonlinearity (black, posterior mean and approximate 95% band) after fitting their gain and baseline. Grey regions indicate stimulus values where the measured nonlinearity is poorly constrained. **(A)** Gaussian prior with its width *σ* varied at *p* = 5.75. **(B)** Standard normal prior with *p* varied. **(C)** Student-*t* prior with its tail index *ν* varied at *p* = 8.5; the inset shows the small differences among these solutions. **(D)** Student-*t* prior with *ν* = 1.5 and *p* varied. In both prior families, changing *p* alters the predicted nonlinearity more strongly than changing the prior, and the closest match to H1 occurs at a large *p*, far from the infomax limit *p* → 0.

